# Systematic analysis of cell morphodynamics in *C. elegans* early embryogenesis

**DOI:** 10.1101/2022.10.31.514462

**Authors:** Yusuke Azuma, Hatsumi Okada, Shuichi Onami

## Abstract

The invariant cell lineage of *Caenorhabditis elegans* allows unambiguous assignment of the identity for each cell, which offers a unique opportunity to study developmental dynamics such as the timing of cell division, dynamics of gene expression, and cell fate decisions at single-cell resolution. However, little is known about the cell morphodynamics, including the extent to which they are variable between individuals, mainly due to the lack of sufficient amount and quality of quantified data. In this study, we systematically quantified the cell morphodynamics in 52 *C. elegans* embryos from the two-cell stage to mid-gastrulation at high spatiotemporal resolution, 0.5 μm thickness of optical sections, and 30-second intervals of recordings. By comparing the morphodynamics between the embryos, we found high reproducibility of the dynamics. The median correlation coefficient of morphological features, such as volume and surface area, was 0.09, suggesting high reproducibility. Furthermore, the morphodynamics was significantly different from the other cell types in all cells before the onset of gastrulation, indicating high uniqueness of the dynamics. Furthermore, physical cell-cell contacts were quantified to analyze inter-cell relationships, and their reproducibility was calculated between the embryos. The number of contacts formed in every embryo was less than half of all detected contacts because variability in division timings and cell arrangements produced various combinations of brief contacts. In contrast, the area of contacts formed in every embryo occupied 96% of the total area, suggesting the high reproducibility of spatial occupancies and adjacency relationships of the cells. Our data further allowed detailed analyses of specific cellular features. By normalizing the sphericity dynamics, we found a significant increase in sphericity at the end of metaphase in every cell, indicating the universality of the mitotic cell rounding. Concomitant with the rounding, the volume also increased in most but not all cells, suggesting less universality of the mitotic swelling. Therefore, we demonstrate that our data provides a precise understanding of the cellular processes and could potentially be used to gain further insight into developmental mechanisms.

## 1 Introduction

*Caenorhabditis elegans* is one of the best-characterized model organisms to study animal development. Its development proceeds through an invariant cell lineage, namely the stereotypical pattern of cell divisions, and produces an adult hermaphrodite with just 959 somatic cells. The whole-cell lineage was firstly established by Sulston *et al*. by manually tracking cells with differential interference contrast (DIC) microscopy (Sulston et al., 1983). Recent advances in bioimage informatics enabled automated tracing of the cell lineage (Onami et al., 2001; Bao et al., 2006). Typical studies perform 4D (3D time-lapse) imaging of embryos with fluorescently labeled nuclei (typically histone) and computationally identify and trace the lineage of the nuclei (Azuma and Onami, 2013). In addition to the nuclear labeling, the use of reporter genes enables the measurement of the reporter expression dynamics at single-cell resolution. This method provides quantified nuclear dynamics, as well as reporter expression dynamics, with the lineage information. The quantified data have allowed systematic analysis of developmental dynamics, including variability of cell division timings and cell cycle lengths measured in 20 embryos (Bao et al., 2008), reproducibility of cell cycle lengths, division axes, and cell positions measured in 18 embryos (Richards et al., 2013), gene expression dynamics in 127 cells (Murray et al., 2012), and high-dimensional phenotypic analysis of 204 essential genes in 1,368 perturbed embryos (Du et al., 2015). In these studies, a large sample size led to the reliability, and automated analysis ensured the objectiveness of the conclusions.

Cell morphology is also associated with a variety of biological processes. Relationships have been found between cell volume and cell cycle length (Arata et al., 2015), cell volume and strength of the spindle assembly checkpoint (Galli and Morgan, 2016), asymmetric divisions and the local inactivation of actomyosin cortical contractility (Rose and Pierre Gönczy, 2013), and asymmetric divisions and confinement of embryos (Fickentscher and Weiss, 2017). However, there has been little systematic analysis of the morphodynamics of cells due to the lack of sufficient amount and quality of quantified data. Consequently, there is little quantitative knowledge about morphodynamics, including the extent to which they vary between individuals.

Although the cell morphology can be visualized by imaging embryos with fluorescently labeled cell membranes, computational identification of the membranes is more challenging than that of nuclei. This is because nuclei are thick, well-separated spherical structures, whereas cell membranes are thin planar structures that contact each other, forming complicated networks. Despite these difficulties, several methods have recently been developed for membrane segmentation. In 2017, we developed an image processing method called Biologically Constrained Optimization-based cell Membrane Segmentation (BCOMS), which automatically segments cell membranes and extracts morphological features of each cell by solving an objective function under biological constraints (Azuma and Onami, 2017). It uses previously segmented nuclei manually validated as markers, yielding cell segmentations with no missed cells. The performance of BCOMS was evaluated by comparisons with manually created ground truth and between two adjacent time points and was the best among the available methods. After this, two other methods have been developed, namely 3DMMS (Cao et al., 2019) and CShaper (Cao et al., 2020). These methods opened the door to a large-scale systematic analysis of the morphodynamics of cells.

In this study, we quantified and systematically analyzed the morphodynamics of cells in 52 *C. elegans* embryos. For the quantification, we updated the BCOMS and applied an image restoration method employing a deep convolutional neural network as a preprocessing to improve the image quality. As the time interval of the recordings was 30 s, which is shorter than half of the typical *C. elegans* studies (1–1.5 min), morphodynamics can be sufficiently captured, including fast motions such as mitosis. Along with a large number of replicates (52 embryos), it enables us to detect and analyze subtle differences in the dynamics with statistical reliability, which is essential for evaluating its reproducibility or variability across embryos. By taking advantage of these, we demonstrated the availability of the data to precisely understand relevant developmental dynamics.

## 2 Results

### 2.1 Quantification of the morphodynamics of cells

We performed 3D timelapse imaging of 52 embryos, in which the nucleus and cell membrane are labeled with green fluorescent protein (GFP) and mCherry, respectively, for two hours from the two-cell stage. We detected the cell nucleus at each time point using an image processing method developed previously (Azuma and Onami, 2013) and curated detection errors using in-house curation software. Consequently, we prepared error-less positional data for the nucleus in all the 52 embryos, which would be used as inputs to BCOMS. As every cell has a name in the *C. elegans* embryo, based on its ancestry, we assigned the cell names for every detected nucleus by referring to publicly available annotated data (Richards et al., 2013, see Methods).

Although the performance of BCOMS was validated, we upgraded it by adding new functions and preprocessing to improve the quantification quality and acquire additional morphological features, which we call BCOMS2. As preprocessing, we developed an image restoration method based on deep learning (see Methods). The method trains a convolutional neural network based on the U-Net (Ronneberger et al., 2015) using consecutively acquired low- and high-quality images. The trained network can restore newly given low-quality images and reconstruct high-quality images. One of the added functions is the collection of the time lag between nuclear divisions (karyokinesis) and cell divisions (cytokinesis). In this period, the two divided nuclei are used as markers for membrane segmentation, although the cell was before the division, leading to incorrect segmentation. The added function collects the time lag by a machine learning-based method (see Methods). The other function is the extraction of cell-cell contacts. These are detected from the cell membrane segmentation results as pairs of cells contacting their surfaces. For each pair, BCOMS2 extracts the contact area and duration.

We applied the BCOMS2 pipeline for the 52 embryos and extracted eight morphological features and two features related to cell-cell contact (Supplementary Table 1, a segmentation result in an embryo is shown in Supplementary Video 1). We evaluated the performance by calculating the volume deviation between two adjacent time points, also used in the previous study (Azuma and Onami, 2017). It can be used as an indicator to evaluate the accuracy of the segmentation because the volume is nearly stable between the adjacent time points under the imaging interval of 30 s. The deviation was reduced from 0.065 to 0.060, demonstrating an improvement in the performance of BCOMS2 by the image restoration method.

The developmental rates can vary even if the imaging conditions are consistent (Schnabel et al., 1997; Bao et al., 2008). The final developmental stages were different between the embryos for the same duration of the recordings. Indeed, the ratio between the fastest and slowest rates was 1.2 in our data. As a result, the number of cells at the final time point varied from 51 to 96 cells, and the number of cells that completed their cell cycle varied from 49 to 119 (Supplementary Figure 1). Of the 52 embryos, 32 embryos exceeded the 85-cell stage and contained at least 76 cell types completing their cell cycle. We used the data from the 52 or 32 embryos for the following analyses.

### 2.2 Reproduction of the previous studies

Some morphological features have already been analyzed in *C. elegans* embryos by others. We picked up two such studies and examined whether our data was consistent with their results. The first study measured volume asymmetry between daughter cells in all cell divisions until the onset of gastrulation (Fickentscher and Weiss, 2017). P lineage cells showed markedly different volume ratios, and divisions of ABar, EMS, MSa, MSp, Ca, and Cp were also significantly asymmetric. In contrast, E, MS, and C underwent almost perfectly symmetrical divisions.

We applied the same scheme and criterion (see Methods) and evaluated the asymmetry of the 27 cell divisions in 52 embryos (Figure 1A). A significant asymmetry was found in all P lineage cells, ABar, MSa, MSp, and Cp, all of which were also detected in the previous study. Especially, P lineage cells showed markedly different volume ratios, and the order of degree of asymmetry was consistent with the previous study. The divisions were almost equal in size in MSa, MSp, and C, congruous with the previous findings. In contrast, the asymmetry was not significant in EMS and Ca cells, which divided asymmetrically in the previous study. Altogether, our results showed agreements with the previous study in 93% (25/27) of divisions.

**Figure 1.**
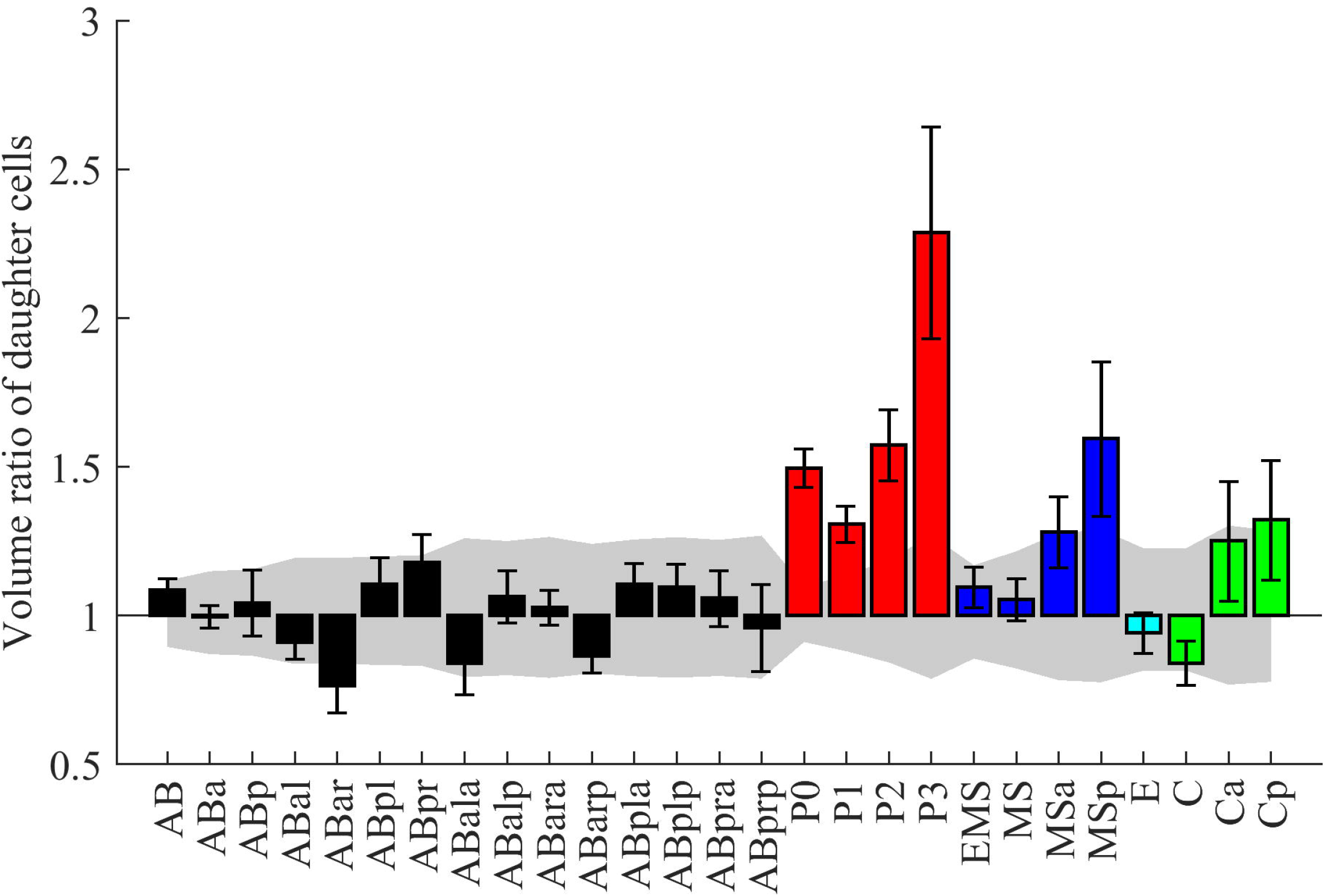
Reproduction of previous studies. (A) Volume ratio of daughter cells emerging from the named mother cell. The design of the graph follows (Fickentscher and Weiss, 2017). The medians of the 52 embryos are shown with error bars indicating standard deviations. Colors indicate cell lineages. The gray region indicates the volume-dependent level of uncertainty (see Methods for details). (B) A double logarithmic plot shows the relationship between cell cycle duration and cell volume for the cells in each lineage. The design of the graph follows (Arata et al., 2015). The data were fitted to the formula, y = a + bx, by the linear least-squares method. The fitting result for the power-law model and coefficient of determination (R^2^) are shown in each graph.

Although their analysis was limited to the divisions until the onset of gastrulation, our data permits the analysis beyond gastrulation. We applied the same analysis for 79 divisions, including the previous 27, in 35 embryos (Supplementary Figure 2). Surprisingly, asymmetry was more significant in Caa than in any of the P lineage cells. The median volume ratio was 2.9 in Caa. We manually checked some original images and confirmed that the segmentation results were correct. In addition, we found 12 new asymmetric divisions. These results demonstrate that our data can be used to reproduce the previous study and acquire new knowledge.

The second study quantitatively measured cell volume and cell cycle duration and found power-law relationships between them, except for the E lineage (Arata et al., 2015). The powers were classified into three classes where the 95% confidence intervals overlapped. The C and P lineages were highly size-correlated (the power was 0.41), AB and MS lineages were moderately size-correlated (the power was 0.27), and E and D lineages were potentially a size-non-correlated class (the power was less than 0.13).

Following their method, we fitted three different models (Gaussian, exponential, and power-law) to the distribution of cell cycle duration vs. cell volume in each cell lineage. The χ2-value in the model fitting was the smallest (except for the E and D lineage) when the data were fitted by the power-law model. The powers were 0.41 in the C lineage, which was classified as the highly size-correlate class, 0.31 and 0.29 in AB and MS lineages, respectively, which were classified as the moderately size-correlated class, and 0.089 and 0.13 in E and D lineages, respectively, which were classified as the size-non-correlated class. P lineage cells were not analyzed because P1 and P4 cells were not recorded over the complete cell cycle length in our data (Figure 1B). Overall, our results showed a good agreement with the previous study.

Overall, we succeeded in reproducing the results in the past studies, demonstrating the reliability and usefulness of our data.

### 2.3 Reproducibility of cell morphodynamics

To this end, we compared the dynamics of eight morphological features in the 76 cell types that completed the cell cycle among the 32 embryos (the results of ABp are shown in Supplementary Figure 3). We observed that cell-type-specific dynamics of the features were reproducible between the embryos. Meanwhile, we also observed deviations in the dynamics, especially in the temporal direction. One possible source of the deviations is the differences in developmental rates between the embryos. The developmental rates vary even if the imaging conditions are consistent (Schnabel et al., 1997; Bao et al., 2008). Indeed, the ratio between the longest and shortest cell cycle length ranged from 1.14 to 1.40, depending on the cell type. However, we also observed variations in absolute values of the features, especially those that were size-related such as volume and surface area (Supplementary Figure 3). Embryo sizes are known to vary under identical conditions (Moore et al., 2013; Richards et al., 2013; Insley and Shaham, 2018). The ratio between the minimal and maximal volume was 1.42 in our data. To minimize these variabilities, we scaled each segmented embryo linearly to approximate its volume to the average of all the embryos and re-extracted the morphological features. We subsequently equalized the temporal lengths of all the cell cycle (see Methods for details). As a result, we obtained spatiotemporally normalized feature dynamics (Figure 2A). We systematically evaluated the reproducibility of the dynamics. As a measure of reproducibility, we calculated the coefficient of variation (CV) at each time point and averaged it across the cell cycle (Figure 2B). The average CVs across all the cell types ranged from 0.042 to 0.15 depending on the feature, and the median was 0.090. The CV was small for the SA/V (surface area to volume ratio) and sphericity.

**Figure 2.**
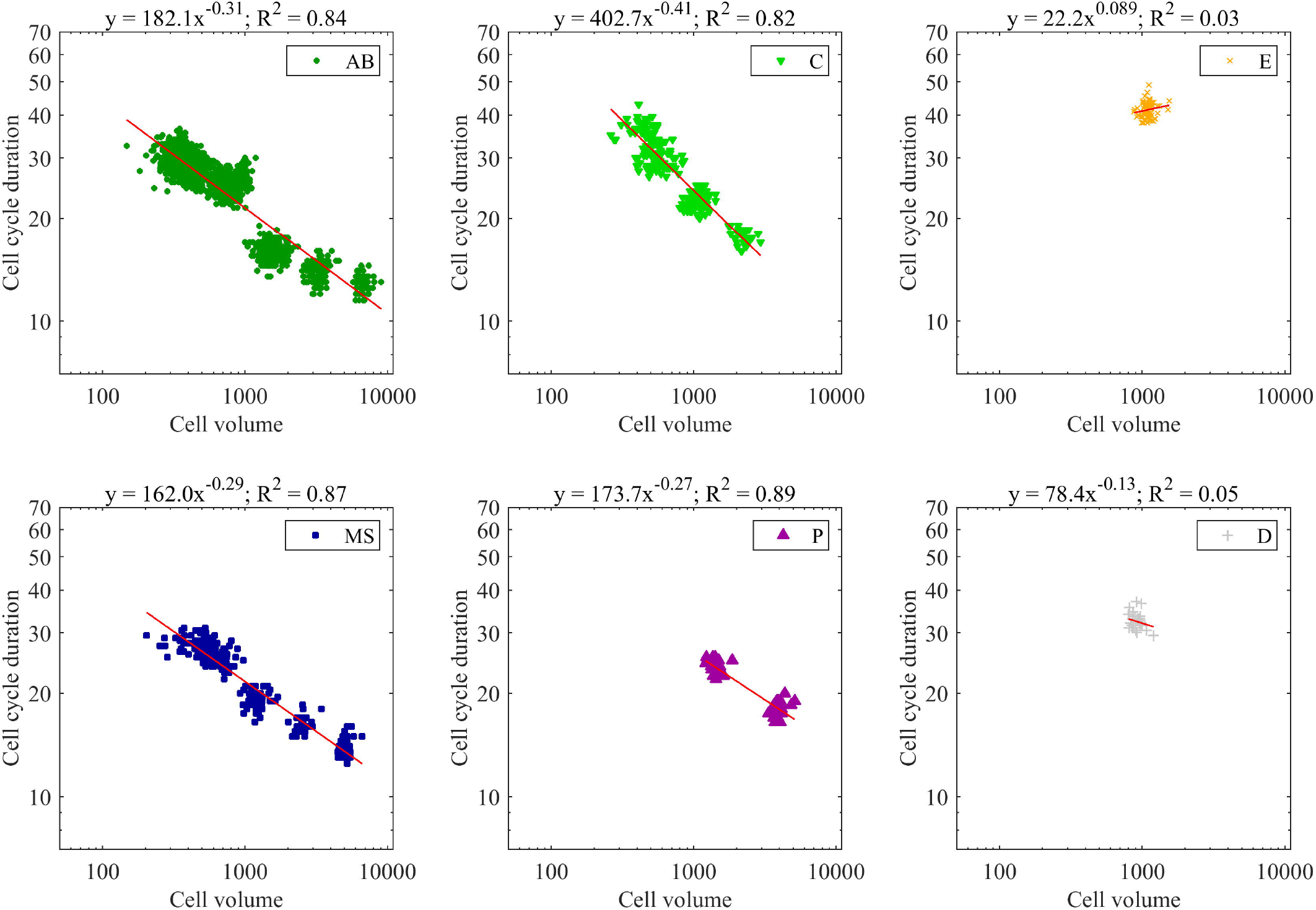
Reproducibility of morphodynamics. (A) Spatiotemporally normalized single-cell feature dynamics in ABp cell. The dynamics of the 52 embryos are shown in different colors. (B) Violin and box plots of the coefficient of variation of the feature dynamics. On each box, the central mark indicates the median, and the bottom and top edges indicate the 25th and 75th percentiles, respectively. The whiskers extend 1.5 times the interquartile range.

### 2.4 The uniqueness of cell morphodynamics

Next, we evaluated the uniqueness of the feature dynamics. If the dynamics are unique for each cell type, they are more similar between the identical cell types than that between the non-identical ones. The similarity can be measured by root mean squared error (RMSE) between any two cells because the temporal length of the dynamics of each feature was equalized. We measured the RMSEs for all pairwise combinations of the 76 types of cells in the 32 embryos; thus, _76*32_C_2_ = 2,956,096 pairs. For each cell, we compared the RMSEs measured with the same cell type in different embryos and the RMSEs measured with different cell types in different embryos. We performed two-sample t-tests (p<0.05) and calculated the number of distinguishable cell types (Figure 3A). The distinguishable cell types were more in volume, surface area, SA/V, long axis, middle axis, and short axis. The SA/V showed higher reproducibility (smaller CV) and higher discrimination power, indicating that it is the optimal feature for distinguishing and identifying each cell type. Indeed, a wide range of bacterial species show a robust SA/V and is suggested to be the parameter the cells monitor (Harris and Theriot, 2018). The cell types distinguishable from others ranged from 6.6% to 41%, depending on the feature. Notably, the distinguishable features were dependent on the cell types. No feature had more discrimination power than any other features in all cell types. This fact raises the possibility that discrimination power may increase upon defining the distance by combining all the features. Simple summation of the RMSEs of the features is dominated by those with larger absolute values because the RMSE roughly correlates with the absolute values of the features. We normalized each RMSE by dividing it by the average of the compared dynamics and summed it across all features. Using this integral RMSE, the cell types distinguishable from any other cell types increased to 68% (Figure 3B). In addition, all cell types could be distinguished from 90% of the others. All cell types before the onset of gastrulation were distinguishable from any other cell types, except for P4. P4 was not included in the analysis because it did not complete the cell cycle in the 32 embryos. When we refer to a cell type by its generation using the lineage name and division times from the founder cell, such as AB2 for ABal, ABar, ABpa, and ABpr, the pairs with non-significant differences contained only AB4, AB5, MS2, and C2, which were born at later developmental stages. We visualized the similarity relationships with the uniform manifold approximation and projection (UMAP) (McInnes et al., 2018; Becht et al., 2019) by using the RMSEs as the distance matrix (Figure 3C). We observed approximately one continuous trajectory where the cells were in order of birth timing (Supplementary Figure 4). The different cell types were separated well in earlier generations and were increasingly mixed along with the progression of embryogenesis.

**Figure 3.**
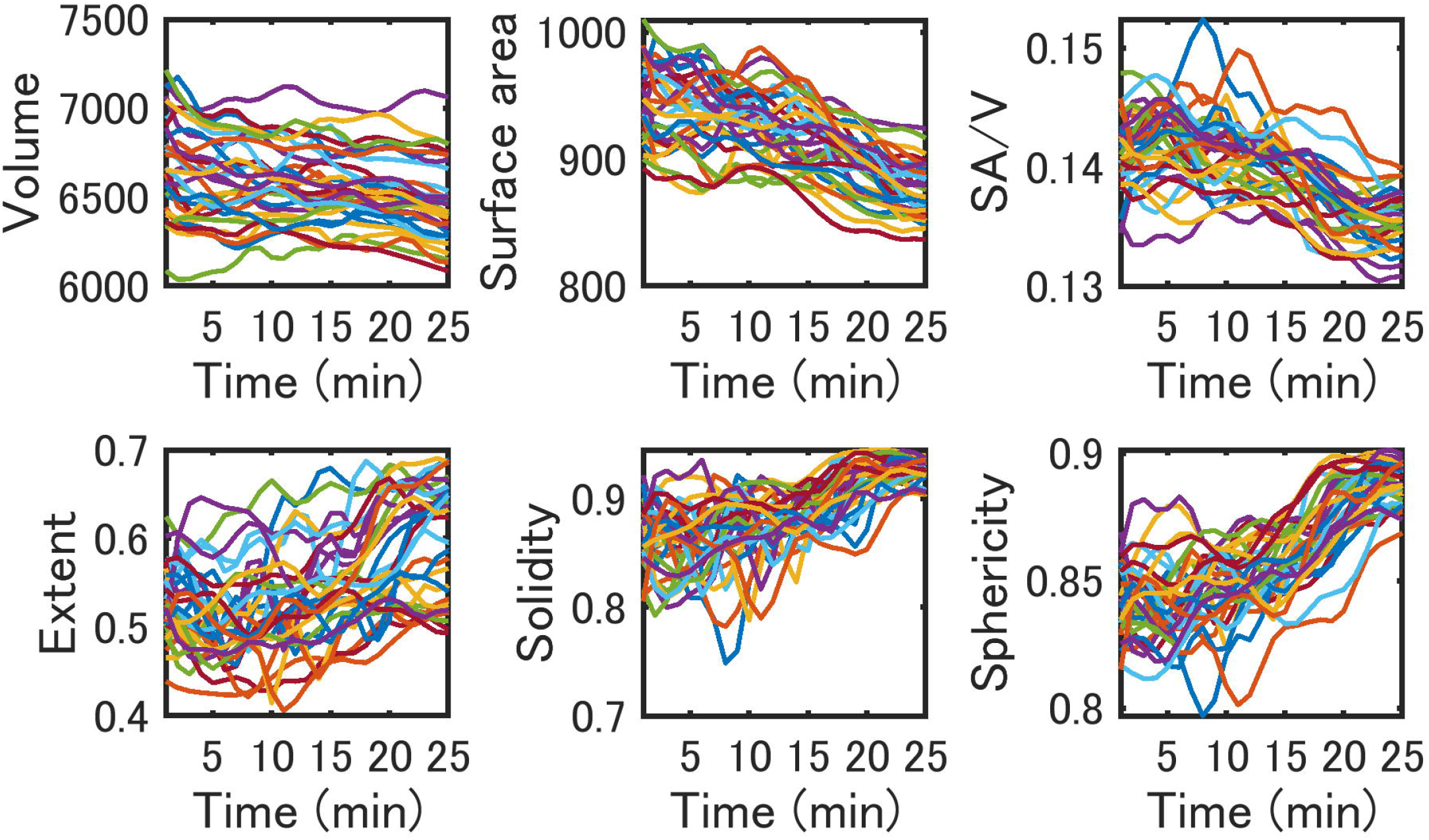
The uniqueness of morphodynamics. (A) Beeswarm boxplots of the cell types that the feature dynamics can distinguish. (B) Frequency of the cell types that the integral feature dynamics can distinguish. (C) UMAP projection of the 76 types of cells in the 32 embryos by using the integral RMSDs as the distance matrix. Colors indicate the cell types. Labels indicate the generation names of the cells.

### 2.5 Reproducibility of cell-cell contacts

Next, we evaluated the reproducibility of the inter-cell relationships, specifically, physical cell-cell contacts across the embryos. The cell pairs forming cell-cell contacts can be automatically detected from the cell membrane segmentation results as those pairs with their surfaces in contact. However, areas of contact as well as the period of contact may range widely. To distinguish each contact by the amount of contact, we computed the integral area by summing the contact area across the cell cycle. The integral areas were determined for all pairwise combinations of the 47 types of cells in the 52 embryos. The cells were selected as those completing the cell cycle in all 52 embryos to evaluate the reproducibility as precisely as possible, because segmentation errors in either of the cell pairs cause false detections. We found a biphasic relationship between the integral area and reproducibility (Figure 4A). In smaller integral areas (below 1,000 μm^2^), the relationship was correlative (r = 0.61). Above 1,000 μm^2^, most contacts were perfectly reproducible across all 52 embryos. Only two contacts (ABplap and ABalpp; ABplap and ABplpp) were imperfectly reproducible. Although ABplap is related to both contacts, these contacts were lost in one different embryo. We manually checked the images of these contacts and found that they were lost in the embryos throughout their cell cycle (Supplementary Videos 2 and 3). The cell arrangements differed in these embryos from those in the other embryos. Moreover, all embryos hatched, further affirming that these contacts are not essential for embryogenesis.

**Figure 4.**
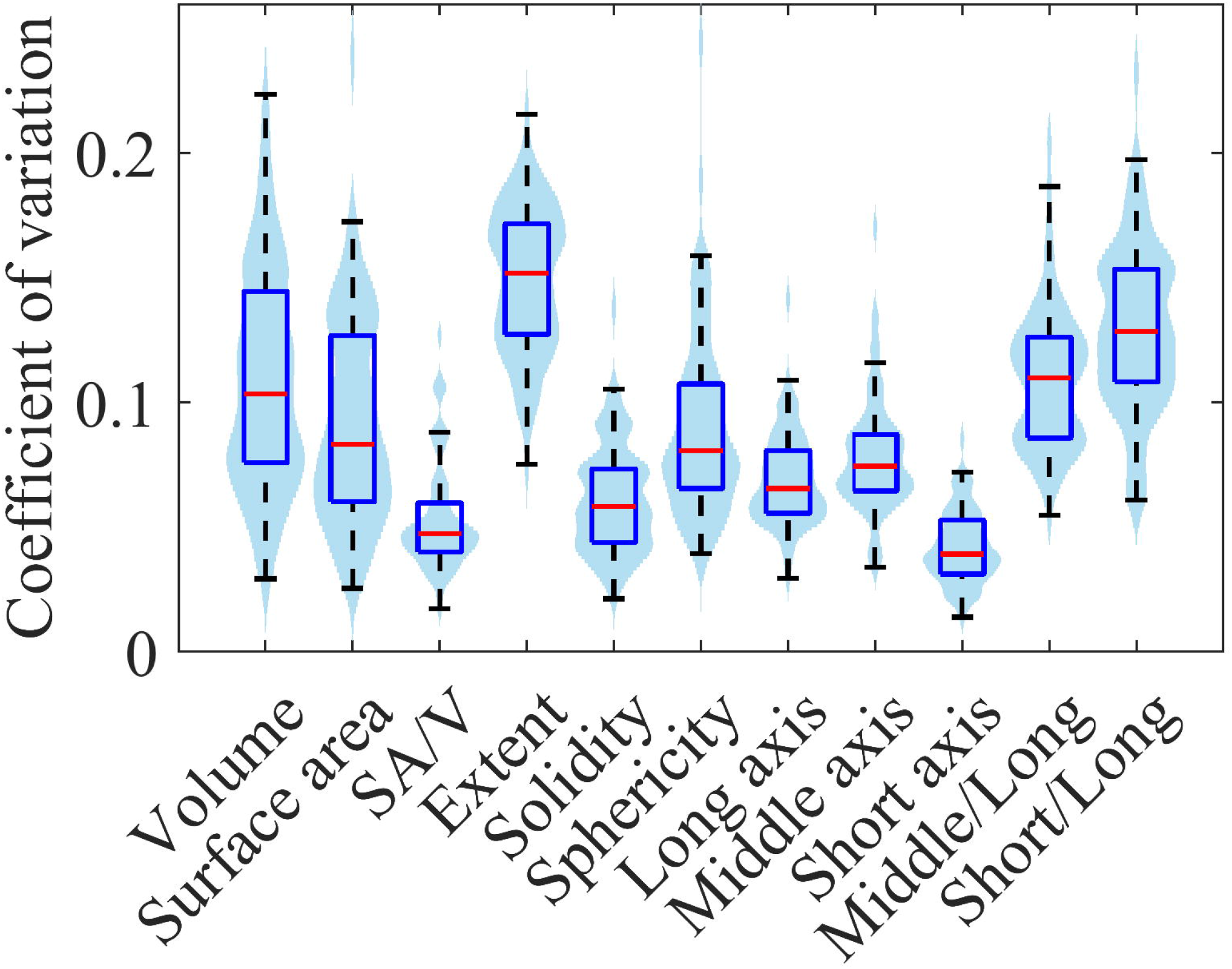
Reproducibility of cell-cell contacts. (A) Relationship between the reproducibility and the integral area of the contacts. The integral area is shown on a logarithmic scale. (B) The reproducibility of the contact over the contact timing. Area indicates the integral area. Contacts with lower reproducibility are observed on both sides, corresponding to cell division timings. (C) Estimated proportions of the three categories of contacts. The number of contacts (left) and integral area (right) in each category are given as a percentage of the total number and total area of all detected contacts, respectively.

As all embryos hatch, the cell-cell contacts mediating the cell-cell interactions should be formed in every embryo. To check this, we selected well-known cell-cell interactions, the five rounds of Notch signaling and two of Wnt signaling (Supplementary Table 2). The cells related to the 4th Notch signaling completed the cell cycle in only 32 embryos. In the cells related to the 5th Notch signaling, parts of the cell cycle were recorded in only two embryos. The reproducibility was evaluated for the limited samples in these cells. All cell pairs showed perfect reproducibility (Supplementary Table 2, Supplementary Figure 5), demonstrating the reliability of our data for contact analysis.

The finding of contacts formed in only one embryo (Supplementary Videos 2 and 3) raises the question of whether more such variable contacts are formed. Most (92%) of the contacts were imperfectly reproducible in those with a less integral area (<1,000 μm^2^). They may include the variable contacts and the false positives and negatives caused by segmentation errors. Among the variable contacts, one expected type is that involving dividing cells. If two adjacent cells divide simultaneously, contacts are formed between mother cells and between daughter cells but are not formed between a mother cell and a daughter cell. If the timings are slightly different, brief contacts are formed between a latter-dividing mother cell and a former-dividing daughter cell(s) in addition to between the mother cells and between the daughter cells (Supplementary Figure 6). Such events frequently occur during *C. elegans* embryogenesis because the division timing is approximately synchronized in cells of each lineage (Sulston et al., 1983) and slightly varies between embryos (Richards et al., 2013). Whether contact is formed between dividing cells can be detected based on when the contact is formed in the cell cycle. The timing can be represented as a single value by calculating the middle time point of the period during which the contact is formed. The values are comparable with each other by normalization to their cell cycle length, which we call “contact timing” (Supplementary Figure 7). The contact timing is 0.5 if the contact is formed throughout its cell cycle. The contact timing is around zero or one if the contact is formed only at the beginning or end of the cell cycle, respectively. Note that two contact timings are calculated for a contacting cell pair, and thus for each of the contacting cells. We computed the contact timings for every contact (Figure 4B). Imperfectly reproducible contacts were observed more at the beginning (0–0.05) and end (0.95–1) of the cell cycle. Perfectly reproducible contacts were not observed at these periods. In contrast, 45 % of the detections were perfectly reproducible at the remaining (0.05–0.95) part of the cell cycle. If the variability is caused by differences in division timings, it is expected to disappear by virtually shifting the division timings back and forth. We realized this by including contacts of daughter cells into contacts of the mother cells. As a result, we observed a decrease in imperfectly reproducible contacts and an increase in perfectly reproducible contacts at the beginning (0–0.05) and end (0.95–1) of the cell cycle (Supplementary Figure 8A). The proportion of imperfectly reproducible contacts decreased to 26% from 100% for these periods (Supplementary Figure 8B), suggesting that the majority of the variability was caused by the variability in division timings.

The remaining variable contacts were caused by false detections or variability in cell arrangements. We manually checked each proportion by selecting 10% of the contacts (23 from 230, Supplementary Table 3). More than half (56.5%, 13/23) were caused by variability in cell arrangements, and false detections caused the others. The estimated total number of variable cell arrangements (130) was nearly one-third (32.8%) of all the cell-cell contacts, while the proportion of the integral area was very less (0.12%).

Using the estimation, we classified all contacts into three categories: perfectly reproducible, variability in contact timing, and variability in cell arrangement, and calculated the proportions of the categories in number and integral area of cell-cell contacts (Figure 4C). The number of perfectly reproducible contacts was not more than half of all contacts. In contrast, the integral area of such contacts was the highest in proportion. However, the variable contacts were more than half in proportion, while the corresponding integral area was small. These results suggest that the spatial occupancy of each cell is highly reproducible, while slight variability in division timings and cell arrangements produce various combinations of brief contacts.

### 2.6 Mitotic cell rounding

Finally, we examined whether our data could be used for elucidating specific developmental processes such as mitotic cell rounding. To clarify this, we focused on the dynamics of sphericity (Supplementary Table 1). In many animal cells, the rounding begins at prophase and the rounded shape is assumed during metaphase until the onset of anaphase, when the cell is elongated, accompanied by chromosome segregation (Ramkumar and Baum, 2016; Taubenberger et al., 2020). Therefore, if the mitotic rounding occurs in the cells of *C. elegans* embryos, the sphericity is expected to be higher, at least at late metaphase. For detecting this period, we used nuclear segmentation data, which was used as a marker in membrane segmentation in the BCOMS2. The end of metaphase is detected as one time point before the segregation of two chromatids. We detected this timing in each of the 76 cell types that completed the cell cycle among the 32 embryos. The sphericity dynamics were registered at the end of metaphase and averaged at each time point over the 32 embryos and were shown to be reproducible between the embryos (Reproducibility of cell morphodynamics section). We observed rapid increases of the sphericity at late metaphase in most of the cell types (Figure 5A). The increase was sharp without a plateau and the peak was within 1.0 min before the end of metaphase in most of the cell types (74/76, Supplementary Figure 9A). Since the time interval of imaging was 0.5 min, the result suggests that the cells kept rounding until the very end of metaphase. The two cell types, ABa and ABp, were an exception and reached a plateau soon after beginning the mitotic rounding, around -4.5 min (Supplementary Figure 9B). The peaks were located several minutes earlier than the end of metaphase in the two cell types.

**Figure 5.**
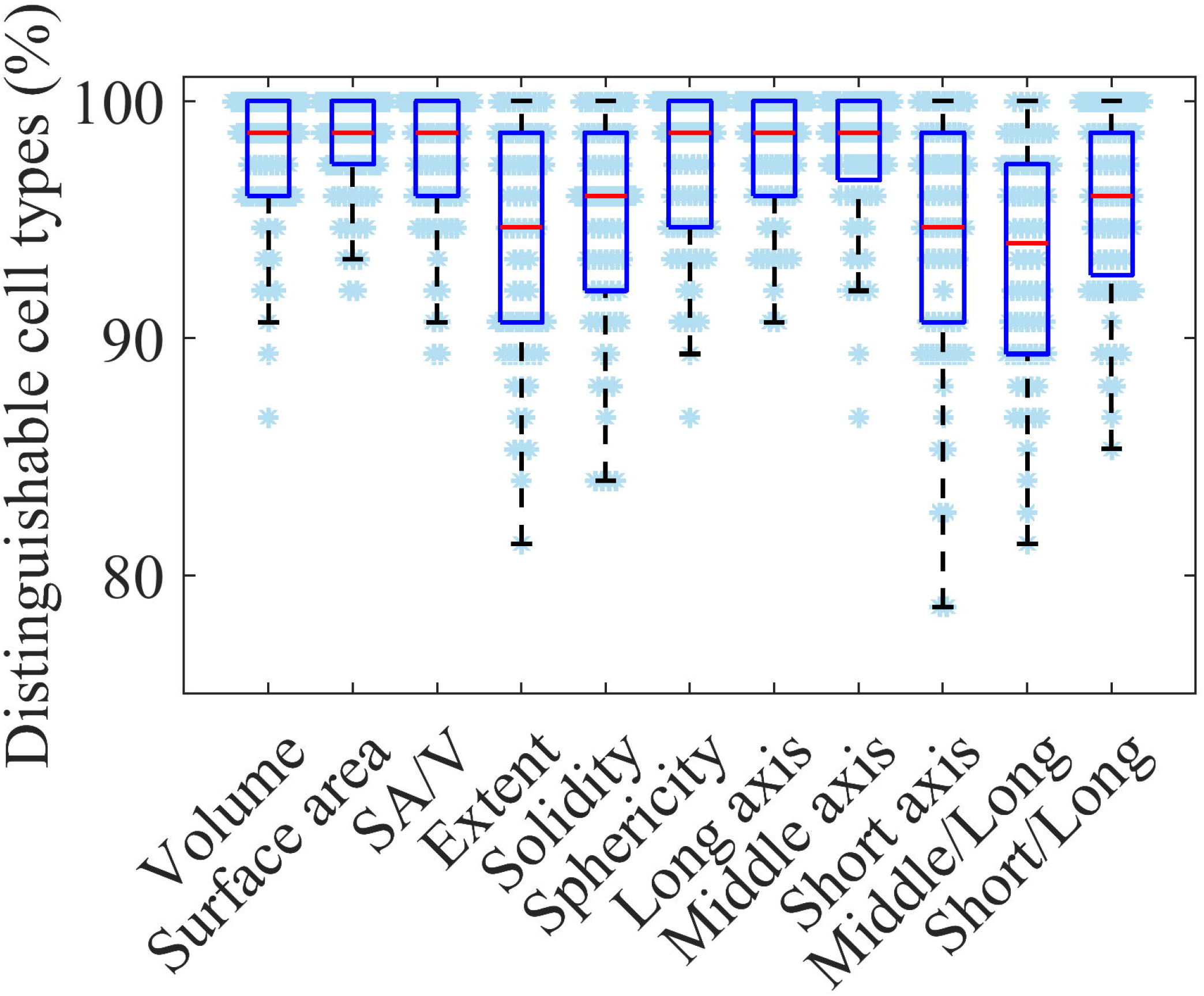
Mitotic rounding and swelling. Sphericity (A) and volume (B) dynamics averaged for the 32 embryos registered at the end of metaphase. The light purple rectangles indicate the initial and end of the estimated duration of the mitotic rounding.

We further averaged the dynamics over all types of cells and found that the sphericity monotonically increased from -7.0 min, when the sphericity is minimum, to 0 min (Supplementary Figure 9C). We defined this period as the duration of the mitotic rounding. We compared sphericity in each cell at the beginning and end of this period. Since the peak positions varied from -1.0 to 0 min, we compared the minimum sphericity during -7.5 to -6.5 min and the maximum sphericity during -1.0 to 0 min. The sphericity significantly increased in all the cell types from 2.0 to 33% (average, 12%) during this period (paired t-test, p<0.05).

The sphericities at the end of metaphase varied from 0.80 to 0.90 (average, 0.85). There was a weak correlation between the sphericity at the initial and the final time points of the mitotic rounding (correlation coefficient, 0.33). However, the correlation was independent of the cell lineage, the developmental stage, and the cell volume (Supplementary Figure 9D-F). Therefore, the final sphericity may depend on other factors, such as the stiffness of the surrounding environment.

### 2.7 Mitotic swelling

We determined whether mitotic swelling occurs in *C. elegans* embryos, which is known to occur concomitant with the mitotic rounding in cultured cells (Son et al., 2015; Zlotek-Zlotkiewicz et al., 2015). We registered the volume dynamics at the final time point of metaphase in the same way as the sphericity (Figure 5B, close-up in Supplementary Figure 10A; embryonic volume was normalized by dividing the volume by the volume at t=0 at each time). Interestingly, the volume dynamics of all types of cells, when averaged, reached a minimum at -7.0 min (Supplementary Figure 10B), which is consistent with the sphericity dynamics (Supplementary Figure 9C). During this period, most types of cells (71/76) significantly swelled from 1.9 to 36% (average, 9.6%; paired t-test, p<0.05).

Among the exceptional five cell types (Supplementary Figure 10C), ABa and ABp showed characteristic sphericity dynamics (see the previous section). Ea and Ep are known to ingress inside the embryo at the beginning of gastrulation (Nance et al., 2005) and have significantly longer cell cycles than the other cells due to the introduction of a Gap phase (Edgar and McGhee, 1988). These characteristics might be related to the disappearance of mitotic swelling. In contrast, we could not find a reasonable explanation for the ABprppp. Although the segmentation accuracy of ABprppp, measured by the volume deviation between the adjacent time points, was the least among the 76 cell types, the same result was observed using only high-accuracy data. Hence, we consider that mitotic swelling does not occur in ABprppp.

We found a correlation in the volume between the initial and final time points of the mitotic swelling (the Pearson correlation coefficient averaged for all cells was 0.73), which was more substantial than that (0.33) in sphericity. A correlation was also reported in the study that measured volume change of cultured cells (Pearson correlation coefficient = 0.53, Son et al., 2015), supporting our results.

Overall, in contrast to the mitotic rounding observed in all cell types, the mitotic swelling was not observed in some cell types, suggesting less universality.

## 3 Discussion

We quantified the morphodynamics of cells in 52 *C. elegans* embryos from the two-cell stage to mid-gastrulation. A large number of replicates allowed reliable evaluation of reproducibility and the uniqueness of dynamics. The high spatiotemporal resolution of the data allowed the detection of subtle and fast changes in cell morphology. We took advantage of these remarkable features and performed distinct types of analyses.

At first, we examined whether our data could reproduce the results in the previous studies to demonstrate the reliability of our data. In comparison with the previous study (Fickentscher and Weiss, 2017) that investigated whether the division is asymmetric or not for 27 divisions during early embryogenesis, our data showed agreements with the study in 25 divisions. Only two divisions were not significantly asymmetric in our data despite being significantly asymmetric in the previous study. The definition of uncertainty partially causes it. The uncertainty (see Methods and Fickentscher and Weiss, 2017) depends on image resolutions. The resolutions were larger in our data in both XY and Z. As a result, the uncertainty in our data was roughly double to triple of the previous study in each cell, which might reduce detection in this study.

The analysis beyond gastrulation found additional asymmetric divisions and markedly different volume ratios between the daughters of Caa. The larger daughter Caaa produces four hypodermal cells, whereas the smaller daughter Caap produces one hypodermal cell, two neurons, and one cell death (Sulston et al., 1983). The high degree of asymmetry may reflect the descendant’s cell death because most cells undergoing cell death are generated as smaller daughters compared with their sisters by asymmetric divisions of their mothers (Hatzold and Conradt, 2008; Conradt et al., 2016). To check this hypothesis, we searched cells whose daughters differed by more than 50% in volume and found seven cells. We traced lineages of their daughters and compared the numbers of descendants undergoing cell death. We found that six of the seven cases followed the hypothesis. Thus, smaller daughters had more descendants undergoing cell death. In the exceptional case, the same numbers of descendants underwent cell death. Therefore, cell death in descendants may be reflected in the high degree of volume asymmetry. The asymmetric division has been suggested to have a functional link to apoptosis (Hatzold and Conradt, 2008). Our results show that smaller daughters had more descendants undergoing cell death. This raises the possibility that asymmetric divisions at several rounds of cell divisions ahead have a functional link to apoptosis. However, the small number of tested samples and an exception require additional investigations to verify this hypothesis.

Our findings were consistent with a previous study (Arata et al., 2015) that found power-law relationships between cell volume and cell cycle duration and classified the founder cell lineages into three classes, except for the P lineage, due to the lack of sufficient data. The reproductions of the past studies demonstrate the reliability of our data. In addition, the extended analysis demonstrates that our data can be used to extend the previous studies.

*C. elegans* has an invariant cell lineage (Sulston et al., 1983), where the pattern of cell divisions and cell fates are reproducible between embryos. Recently, reproducibility was also identified in cell cycle lengths, division axes, and cell positions based on automated nuclear tracking in 18 embryos (Richards et al., 2013). However, reproducibility has not been clarified for cell morphology. Our study showed that the dynamics vary from 4.2 to 15%, and the RMSEs were significantly smaller between identical cells than between non-identical cells by 6.6% to 41%, depending on the feature. Furthermore, the proportion increased to 68% by combining all the features. In addition, all cell types could be distinguished from 90% of the others. These results suggest that the morphodynamics is highly reproducible and unique for each cell.

The analysis benefitted from the high number of replicates in our data. If the number of replicates is half (16 embryos), the proportion of distinguishable cells decreases from 68 to 55%. If the number of replicates is a quarter (eight embryos), the proportion decreases to 37%. Since the developmental rate is different between the embryos (up to 1.2 times), the number of cells at the final time point and the number of cell types completing the cell cycle are different despite starting from the two-cell stage and with the same recording period. Therefore, the more cell types selected, the lesser the number of replicates, and vice versa. To balance these two, we plotted the relationship (Supplementary Figure 1) and selected the 76 cell types in the 32 embryos.

In *C. elegans*, cell-cell interactions play essential roles in embryogenesis. One such interaction, i.e., Notch signaling, plays a significant role in specifying cell fates and tissue morphogenesis (Priess, 2005). It requires cell-cell contacts to transmit the signal (Kopan and Ilagan, 2009). Similarly, Wnt signaling requires cell-cell contacts for signal transmission (Walston et al., 2004). Further, the polarization of blastomeres by restricting PAR proteins also requires sensing the cell-cell contacts by E-cadherin HMR-1(Nance, 2014; Klompstra et al., 2015). However, the reproducibility of the cell-cell contacts has not been clarified. Hence, we evaluated the reproducibility of cell-cell contacts and found a biphasic relationship between the integral area and reproducibility. In smaller integral areas, the relationship was correlative, while in larger integral areas, most of the contacts were perfectly reproducible. The integral area was introduced to minimize the influence of false detections of the contacts, which is inevitable unless the membrane segmentation is perfect, which is almost impossible with current technology. A straightforward way to remove false detection is a threshold-based method. Thus, detections whose contact areas are below the threshold are false positives. However, the detections are highly dependent on the threshold, which is not easy to decide reasonably. Indeed, a previous study that used thresholds for contact area and duration to remove false positive detections suffered from false negatives (Cao et al., 2020). In contrast, our integral area-based approach removes no detections, including false positives. Thus, no false negatives occur. Although the false positives are detected, integral areas are usually small because false detection rarely continues over multiple time points. The integral area-based approach provided us with a new perspective on cell-cell contact.

Evaluation of reproducibility through a simple binary judgment, where all detections were equally contributing irrespective of the integral area, showed that the number of perfectly reproducible contacts was fewer than half of the total contacts, suggesting low reproducibility of cell-cell contacts. In contrast, the evaluation by the integral area showed that the perfectly reproducible contacts occupied most of the total integral area, suggesting high reproducibility of spatial occupancy of the cells. In the Notch signaling pathway, cells need to form physical contacts with more than a certain width and for a certain period to transduce the signal (Kopan and Ilagan, 2009). Our analysis showed that the integral area was higher in contacts mediating the cell-cell interactions (Figure 4A). Hence, the integral area-based classification of the contact may help find new cell-cell interactions, which are expected to have a higher integral area.

Finally, we focused on mitotic rounding and swelling to demonstrate that our data could be used for elucidating specific developmental processes. Mitotic cell rounding is a process by which cells round up to become spherical as they enter mitosis to create space for spindle formation (Ramkumar and Baum, 2016; Taubenberger et al., 2020). It has been commonly observed in vivo during development in many animals, including mouse (Luxenburg et al., 2011), fly (Kondo and Hayashi, 2013; Rosa et al., 2015; Chanet et al., 2017), and zebrafish (Hoijman et al., 2015). However, little is known about *C. elegans*, including the embryonic cells in which it occurs. We used nuclear segmentation data to identify the onset of anaphase as the timing when two chromatids segregate. By registering the sphericity dynamics at the timing, we found that all cells significantly increased their sphericity at the end of metaphase, indicating the universality of the mitotic rounding across embryonic cells. Some cells showed a slight increase (2.0% in the minimum case). Detection by the human eye is nearly impossible for such a slight difference in three-dimensional images, demonstrating the ability of the quantified cell morphology resource.

It had been controversial whether cells increased or decreased their volume during mitosis. In 2015, two studies on the same issue developed distinct methods to precisely measure the volume dynamics of adherent or suspended cells and observed an increase in cell volume (Son et al., 2015; Zlotek-Zlotkiewicz et al., 2015). The increase was observed in cells from a variety of tissues in human and mouse (Zlotek-Zlotkiewicz et al., 2015). However, it is unclear whether the mitotic swelling occurs in vivo, especially in confined environments, including the *C. elegans* embryo, which is enclosed in an eggshell. A previous study found the cell volume to increase concomitantly with mitotic rounding, reach a maximum during metaphase, and decrease shortly before anaphase (Zlotek-Zlotkiewicz et al., 2015). Therefore, if the mitotic swelling occurs in the cells of *C. elegans* embryos, their volume should increase during the mitotic rounding. We found a significant volume increase in most cells (71/76) concomitant with the mitotic rounding. The increase ranged from 1.9 to 36%, depending on the cell, comparable to those measured in a large range of mammalian cell types (Zlotek-Zlotkiewicz et al., 2015). Among the exceptional five cells, we found possibilities that might be associated with the lack of mitotic swelling in the four cells. Further studies are needed to elucidate the mechanism.

## 4 Methods

### 4.1 Imaging

Sample preparation and imaging methods are described in (Azuma and Onami, 2017), except that a time interval of 30 s was used. The number of focal planes differed in each embryo. We recorded 52 embryos for two hours from the two-cell stage. We confirmed that all embryos hatched.

### 4.2 Image restoration

In our study, the spatial and temporal resolutions and the total recording period were set based on a lower laser power and shorter exposure than the optimal settings to avoid photodamage of the biological samples. This caused the signal-to-noise ratio to reduce. We restored the degraded image using a deep learning method we developed similar to the content-aware image restoration method (Weigert et al., 2018), based on U-Net architecture (Ronneberger et al., 2015). The code and detailed network architecture are available at https://github.com/bioimage-informatics/restworm. The network was trained on registered pairs of low- and high-quality images acquired by quickly changing laser power and exposure time. The images were acquired in sparser spatial and temporal resolutions than actual settings to prevent photobleaching and for unbiased sampling throughout development and across optical sections (Supplementary Figure 11). We acquired the images of 10 embryos for network training. Then, we applied the trained network to the images of the 52 embryos.

### 4.3 Cell name assignment

To annotate the detected nuclei, we used previous data as a reference (Richards et al., 2013). A target data that is a set of nuclear coordinates at the final time point was taken. Reference data was selected from the annotated data as a set of nuclear coordinates with the same number of nuclei. If there were multiple such data, all of them were selected one by one. Principal component analysis (PCA) was applied to each target and reference embryo and mapped to each PC coordinate system. Then, the target embryo was rotated around the first PC axis at two-degree intervals and slightly swung along the other PC axes. In each step, the target embryo was scaled to fit in size to the reference embryo by multiplying a scaling factor to equalize the variance of cell positions in each axis and was applied the matching function, which measures the distance for every cell pair between the target and reference, and the distance was used as “cost” for matching them. The minimum cost matching was found using the Hungarian algorithm (Munkres, 1957). The matching was calculated for every step, and the cost was recorded. Finally, the matching with the minimum cost was selected as the annotation for the target data. By using the annotated cells at the final time point, the cells at earlier time points can be successively annotated by tracing back the lineage. If the annotation at the final time point is accurate, all cells can be accurately annotated until the two-cell stage. If it fails on the way, the annotation at the final time point is not accurate. In this case, a set of nuclear coordinates at one time point earlier than the final time point was used as the target data and annotation. The process was repeated until the tracing was successful.

### 4.4 Collection of the time lag between nucleus and cell divisions

We determined the end of cell divisions when the cell membrane completely encloses the cell. We estimated the timing by a support vector machine with eleven features, such as intensity between two divided nuclei in the membrane image (Supplementary Table 4). The training data was manually created for 147 cell divisions in six embryos. The error rate was 1.7% in the 5-fold cross-validation.

### 4.5 Volume ratios of sister cells

The evaluation of cell volume and the significance of asymmetry was done according to the method described previously (Fickentscher and Weiss, 2017). The median volume across the cell cycle was used as the volume. Cell divisions were deemed significantly asymmetric if the volume ratios of their daughters exceeded uncertainty levels, which arise solely from segmentation errors. In short, the uncertainty is calculated as the sum of voxels comprising a layer around each cell.

### 4.6 Cell cycle length normalization

Cell cycle lengths of a given cell vary between the embryos due to differences in developmental rates and fluctuations of division timings. The lengths ranged from 12 to 49.5 min for all the cells in the embryos, and the average was 25.2 min. Therefore, we normalized the temporal lengths of the feature dynamics by spline interpolation to make their length 25 min (50 time points).

### 4.7 Embryonic size normalization

Our membrane segmentation method (BCOMS2) segments embryonic regions before membrane segmentation. Embryo volume was calculated at each time point by summing all the pixels of the embryonic region. Since the embryonic volume changes slightly throughout development, the median volume was used as the volume of the embryo. Then, an average embryonic volume was calculated from all the embryos. Each embryo image and each segmentation result were linearly expanded or shrunk to approximate its volume to the average volume.

### 4.8 Detection of contacting cell pairs

One cell was picked from the cells of an embryo, and its region was three-dimensionally dilated with a structural element of the one-pixel radius (3×3×3). The dilated region overlaps with surrounding cells in case of contact. The number of overlapping pixels was counted for each contacting cell, and the image resolution was multiplied to calculate the contact area. The area was summed up throughout the embryogenesis period and termed as the integral area. Every cell of the embryo was successively picked, and the areas were calculated. Since the integral area was calculated for each contacting cell, two integral areas were calculated in each cell pair. The two areas were averaged and used as the integral area of the cell pair.

## 5 Data Availability Statement

All datasets are available via SSBD:repository (Tohsato et al., 2016) (https://doi.org/10.24631/ssbd.repos.2022.06.236). All the software developed in this study can be available from https://github.com/bioimage-informatics.

## 6 Author Contributions

Y.A. designed the study, developed the software, analyzed the data, wrote the manuscript, and prepared the figures and tables. H.O. performed the experiments. S.O. supervised the study and wrote the manuscript.

## 9 Acknowledgments

The authors thank all members of the Onami laboratory for input and discussion.

## 10 Supplementary Material

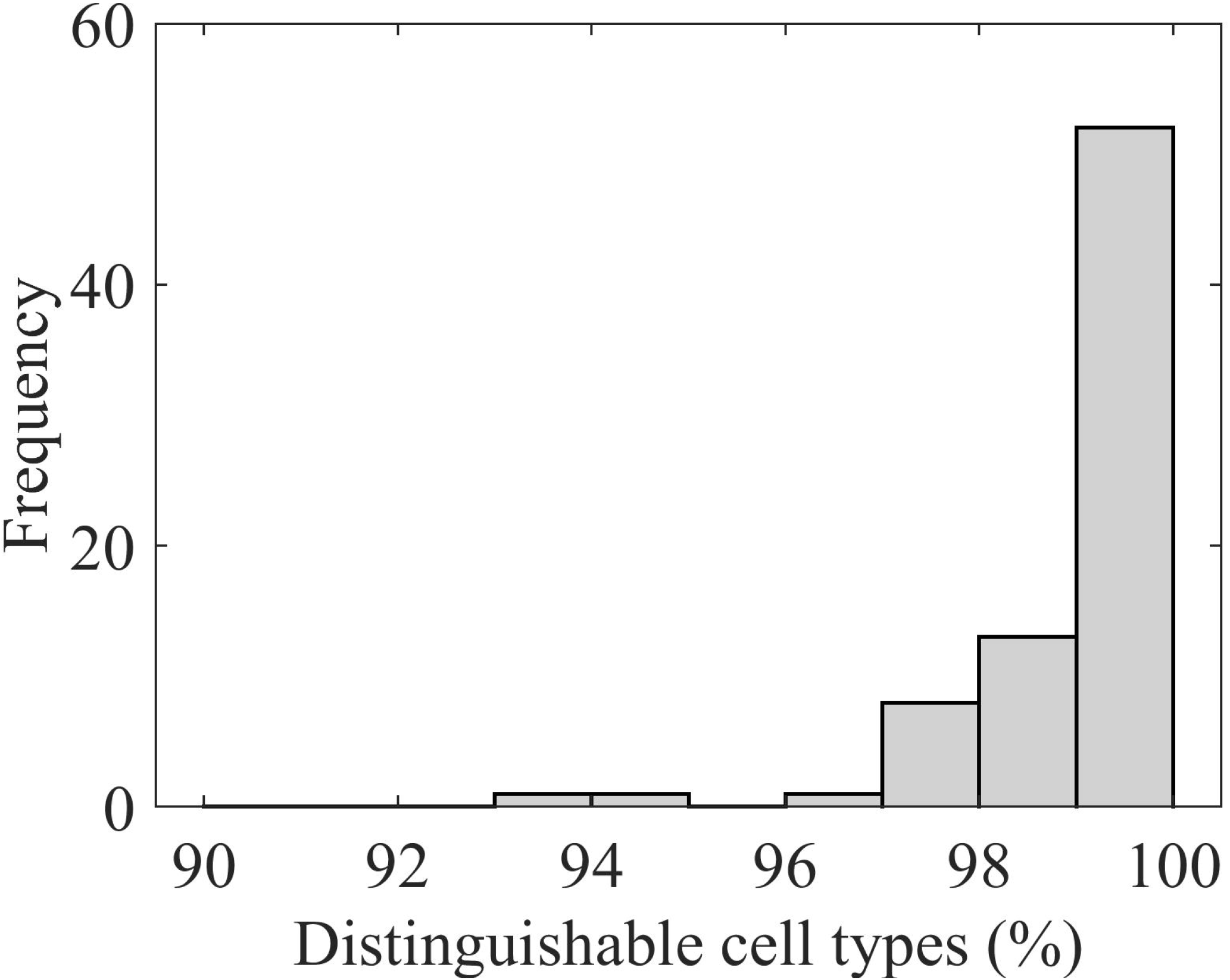

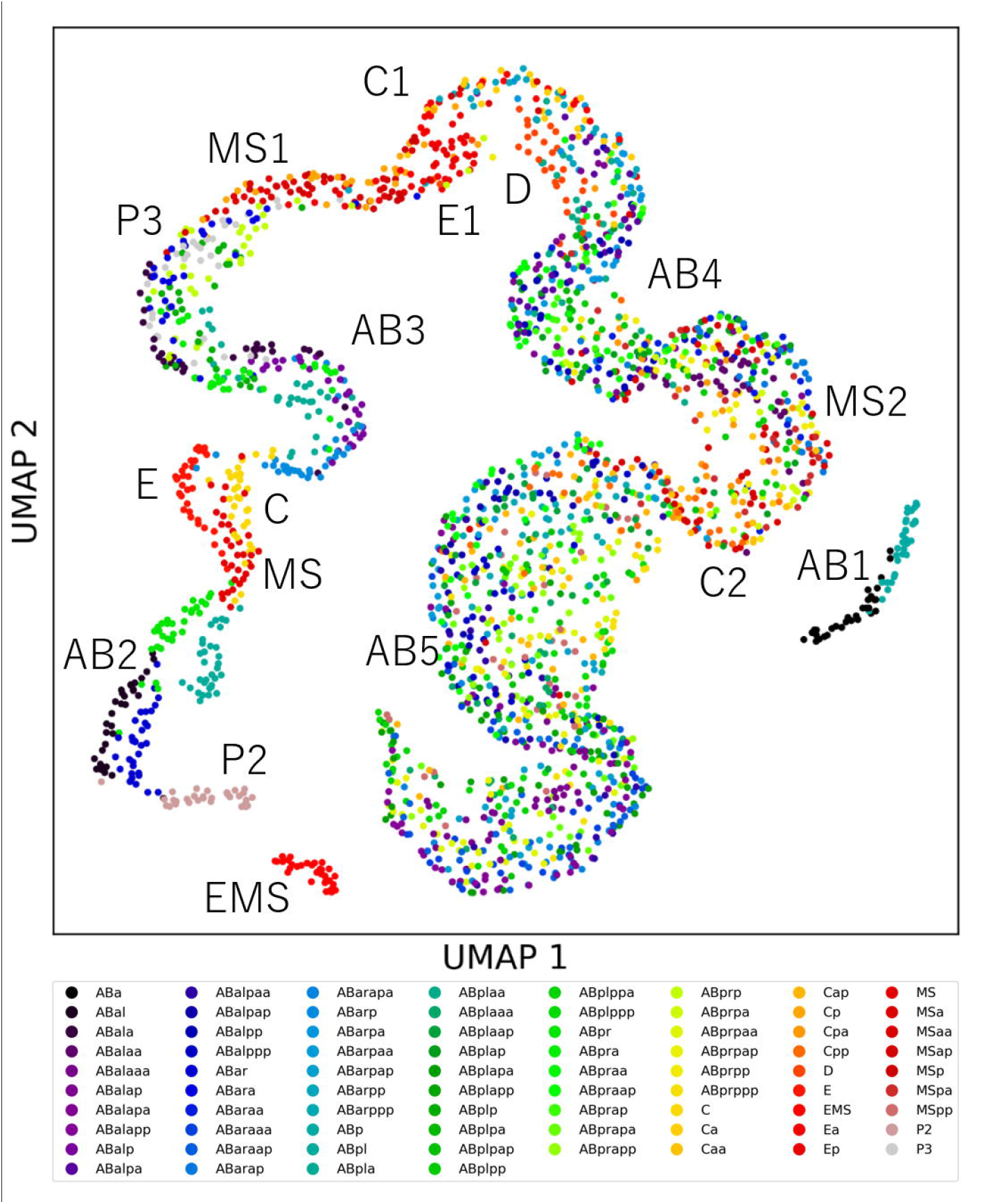

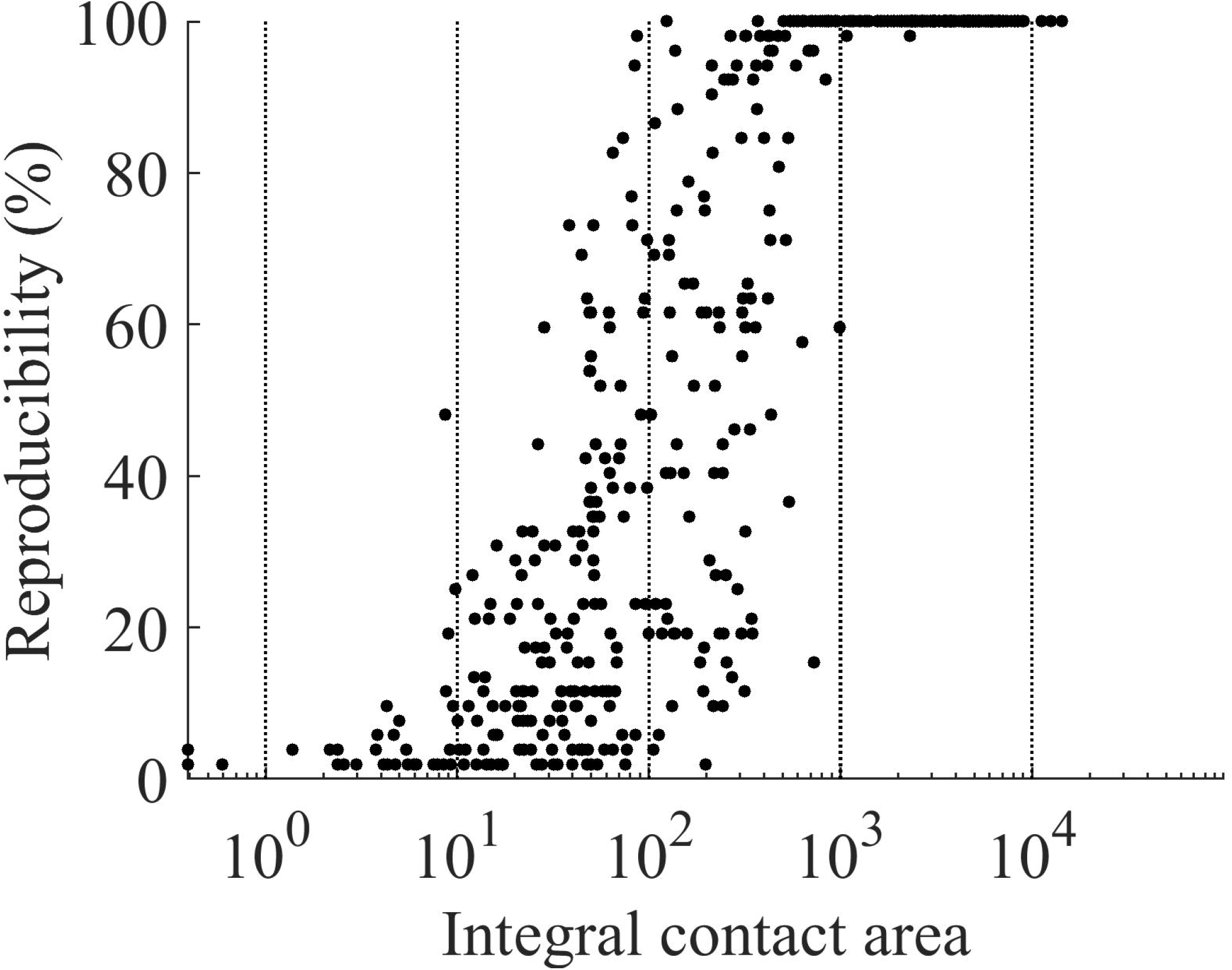

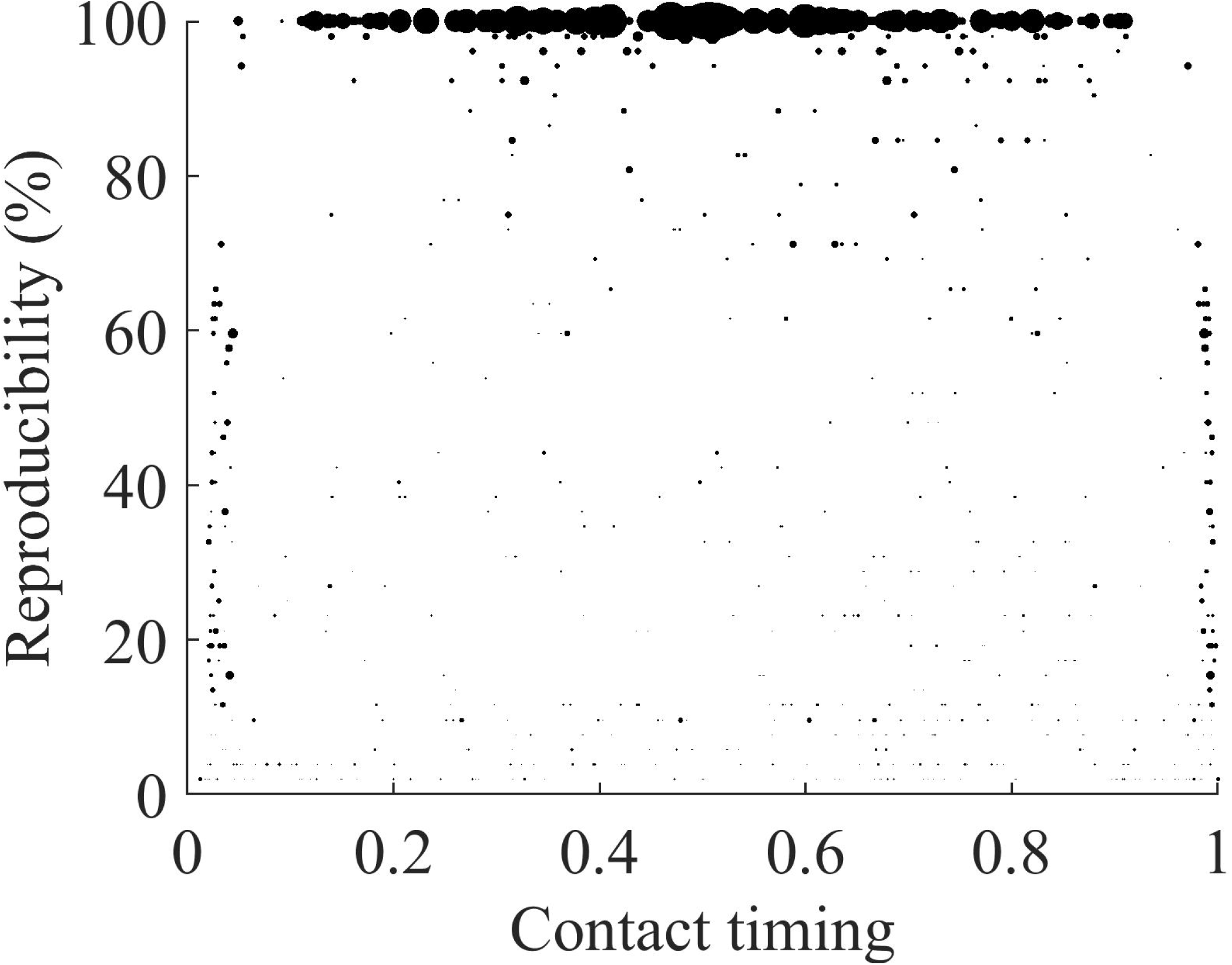

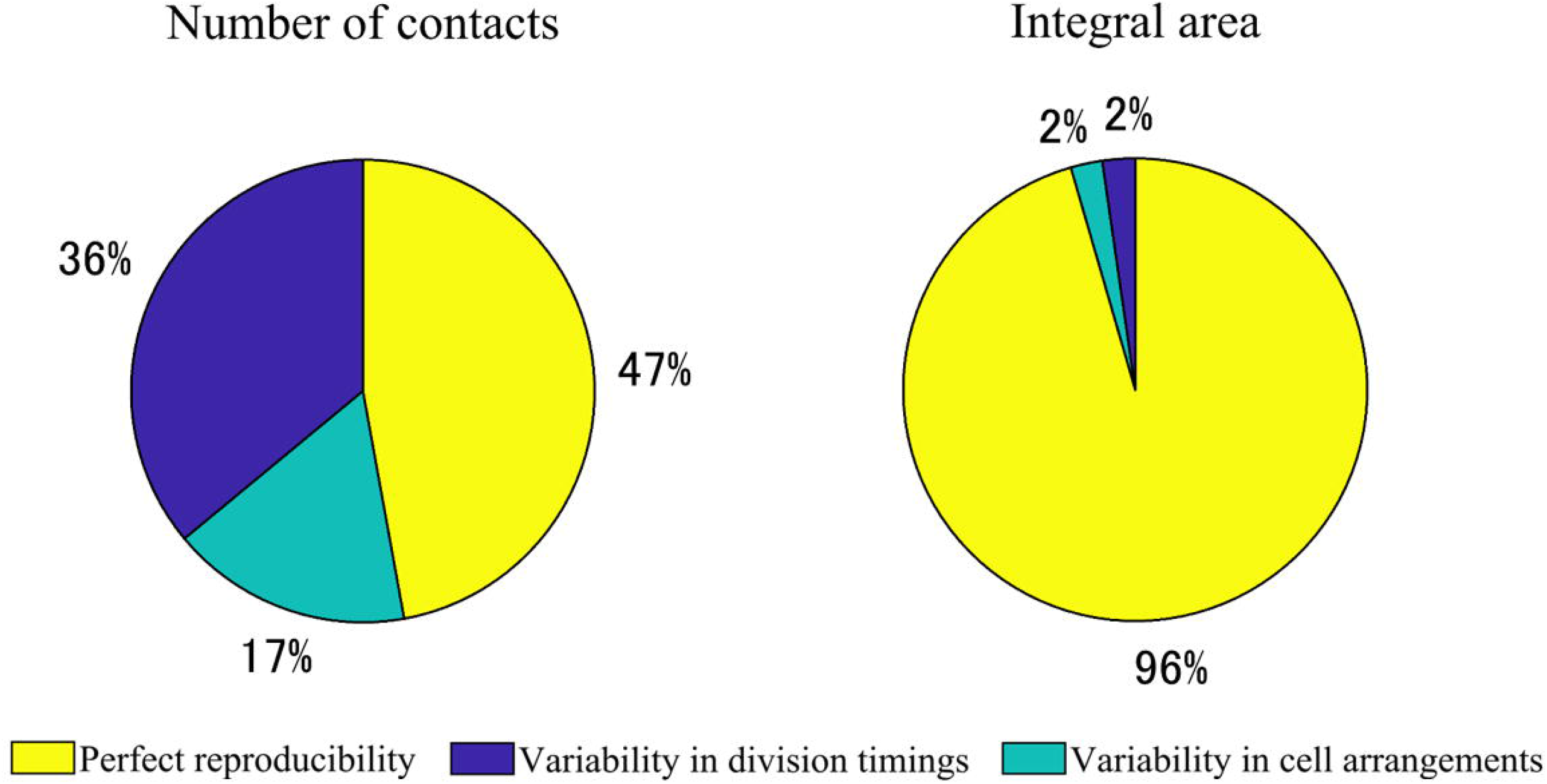

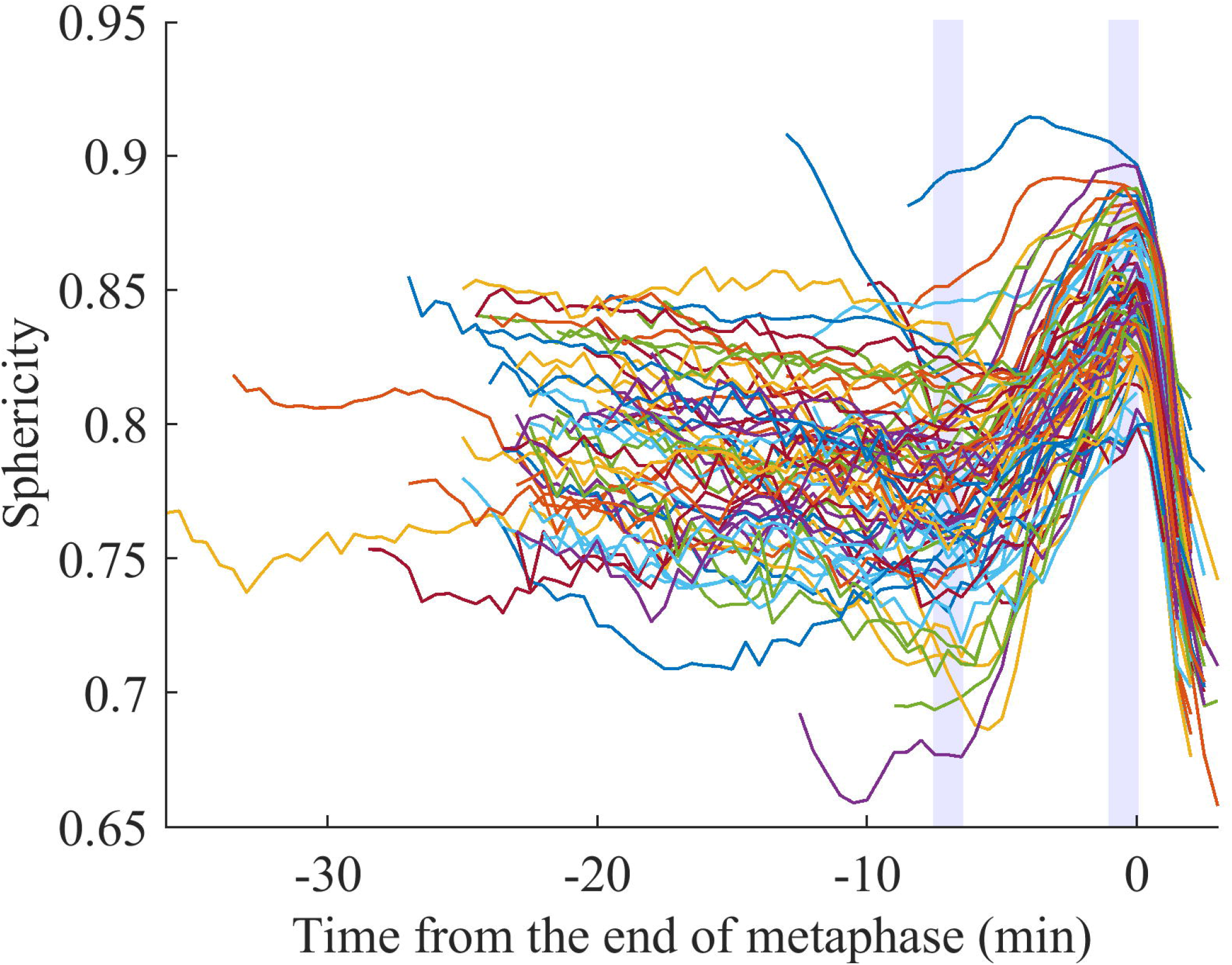

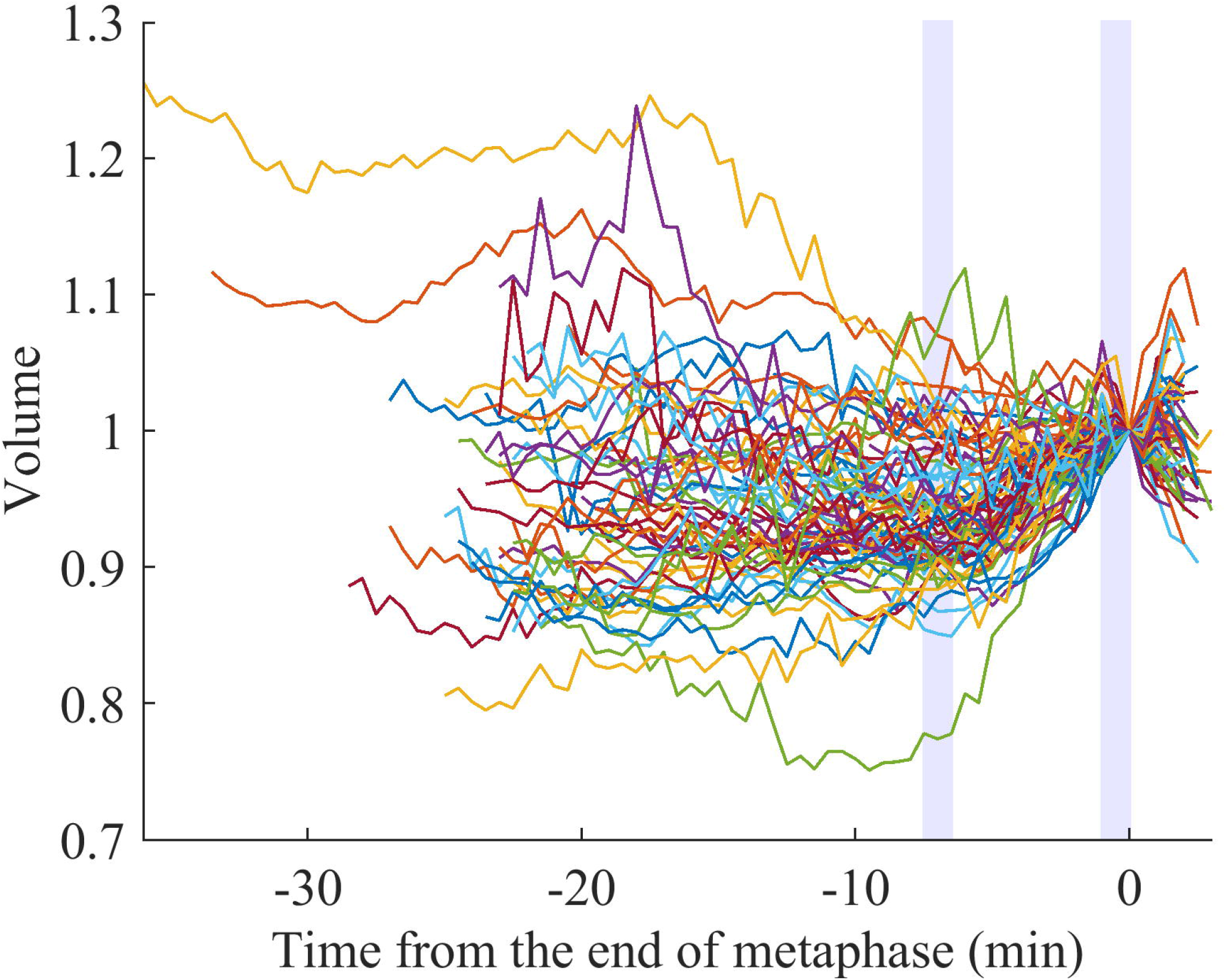

