## Supplementary Material for "Systematic analysis of cell morphodynamics in *C. elegans* early embryogenesis"

### 1 Supplementary Figures and Tables

#### 1.1 Supplementary Figures

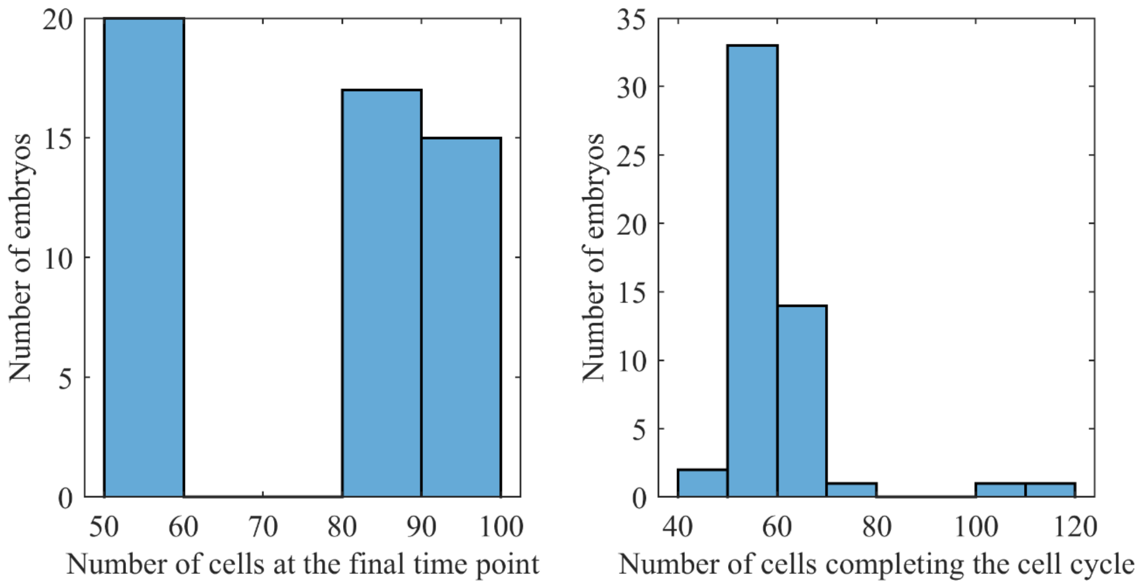


**Supplementary Figure 1.** Statistics of the cells and embryos.

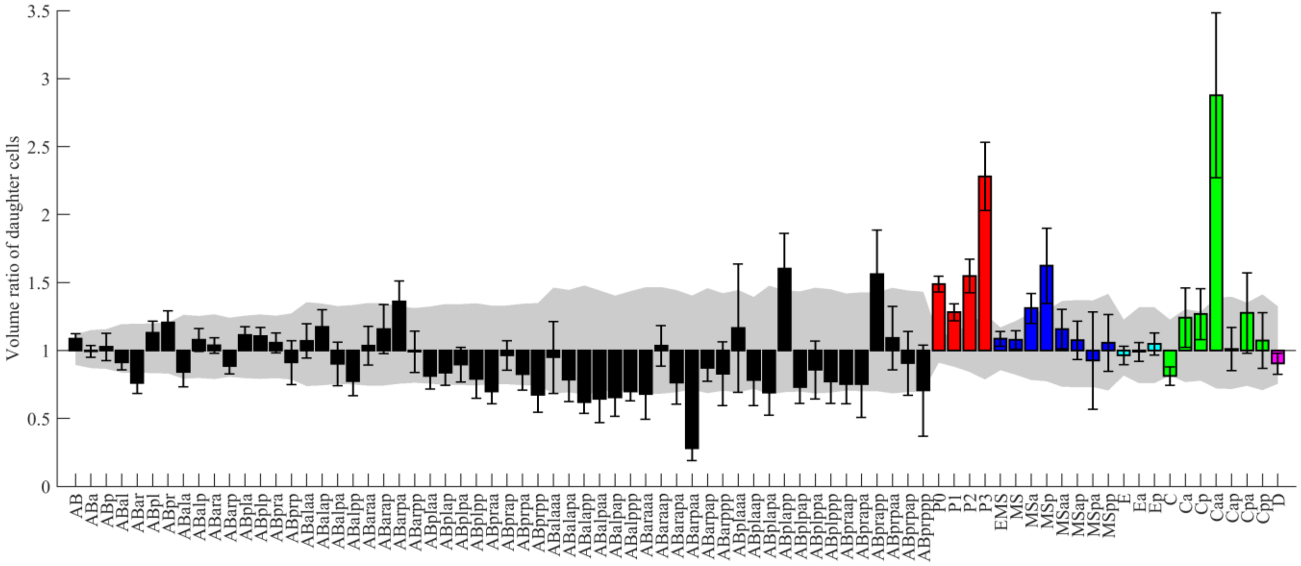


**Supplementary Figure 2.** Volume asymmetry of all cell divisions in our data.

Volume ratios of daughter cells emerging from the named mother cell. The design of the graph is the same as in Figure 1A. The medians of the 52 embryos are shown with error bars indicating standard deviations. Colors indicate cell lineages. The gray region indicates the volume-dependent level of uncertainty (see Methods for details).

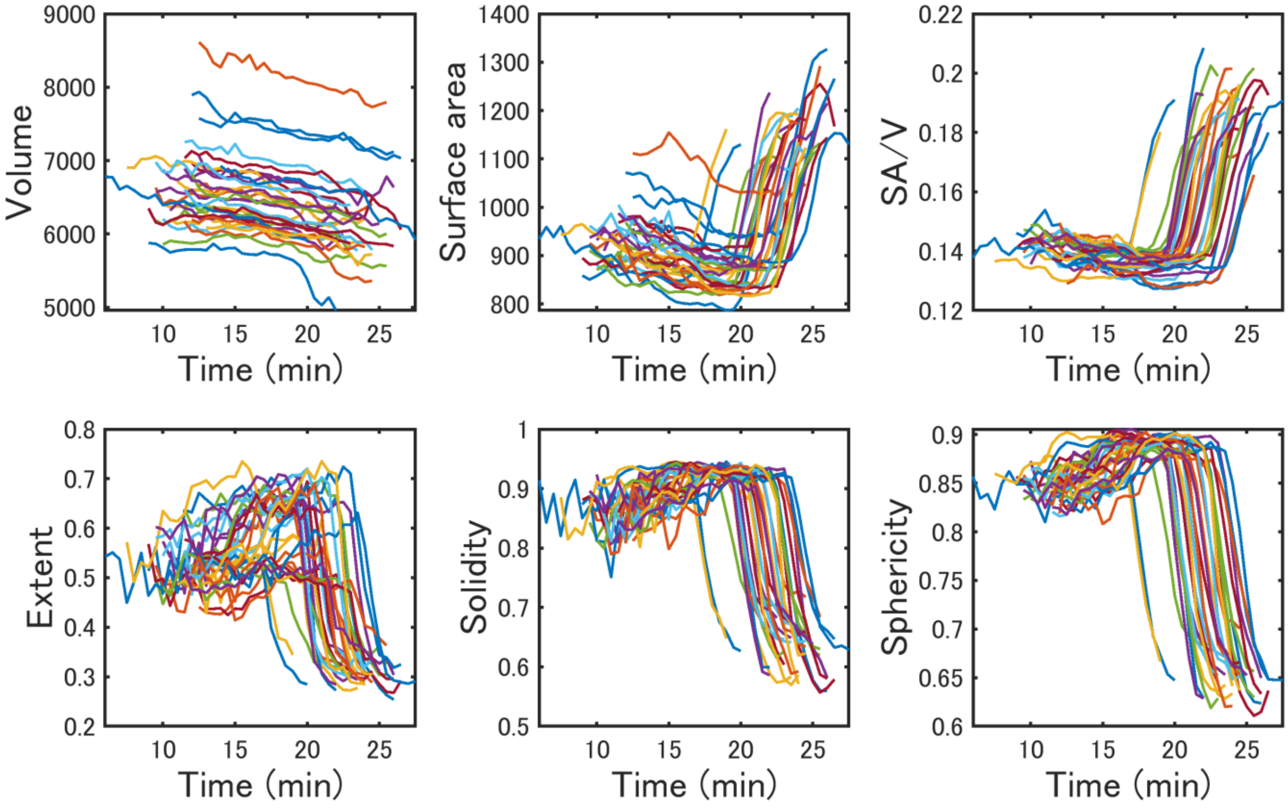


**Supplementary Figure 3.** Dynamics of single-cell features in ABp cell.

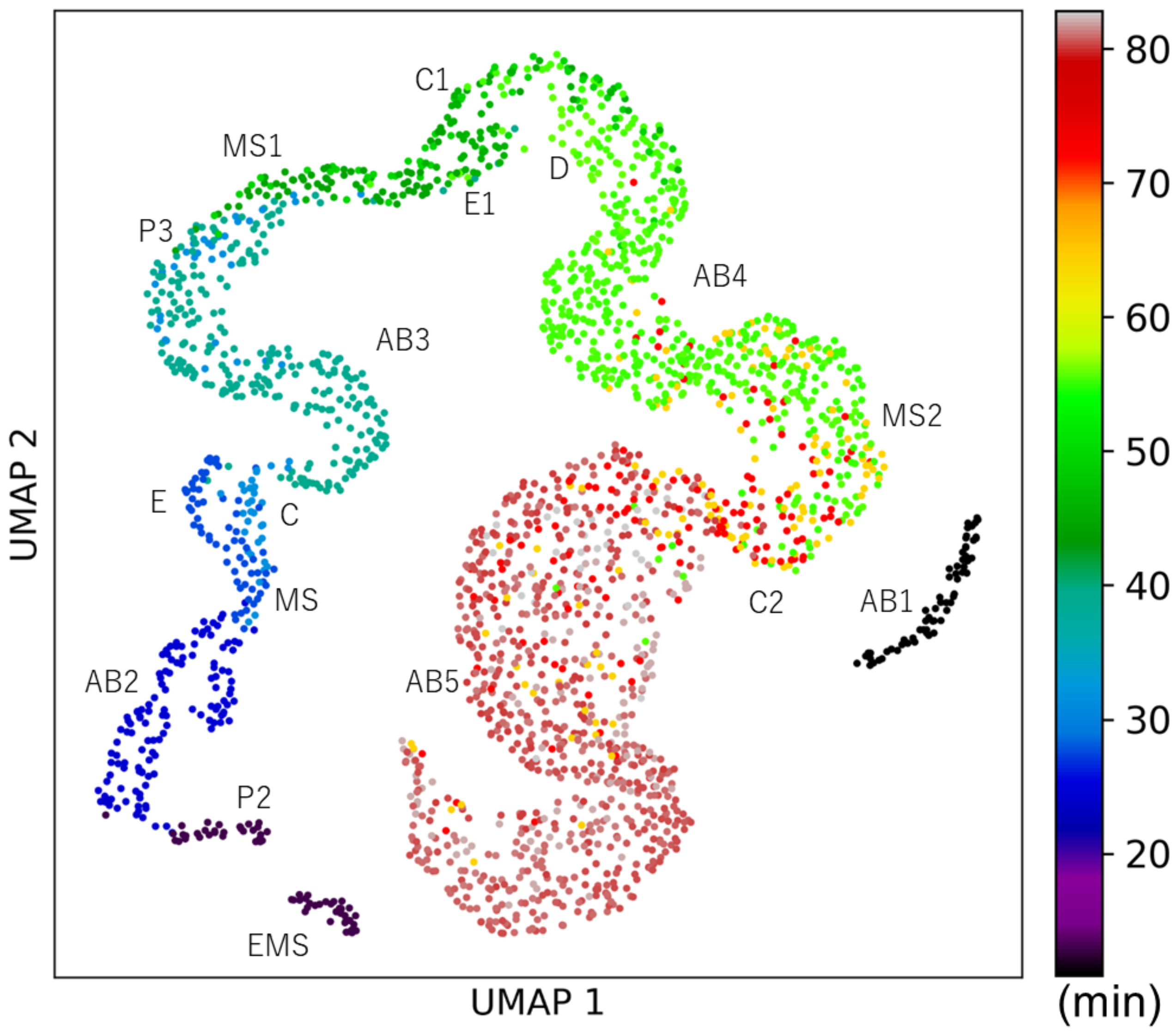


**Supplementary Figure 4.** Umap projection colored by birth time.

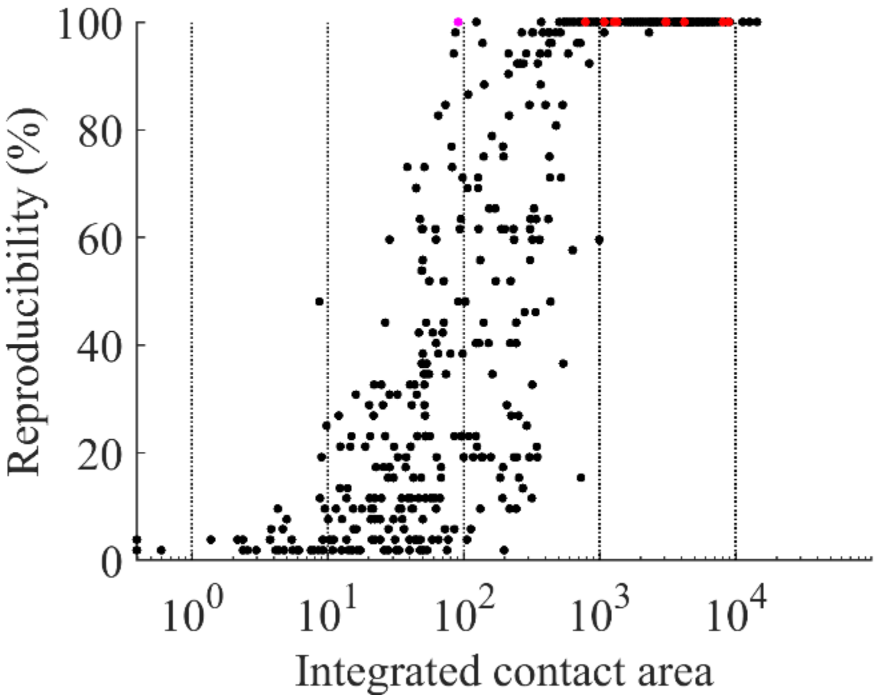


**Supplementary Figure 5.** Reproducibility of cell-cell contacts mediating the cell-cell interactions.

Relationship between the reproducibility and the integral area of the contact. The integral area is shown on a logarithmic scale. The contacts mediating the cell-cell interactions are shown in red. The contact mediating the 5th Notch signaling is added (colored magenta), although its cell cycle does not complete in any of the embryos.

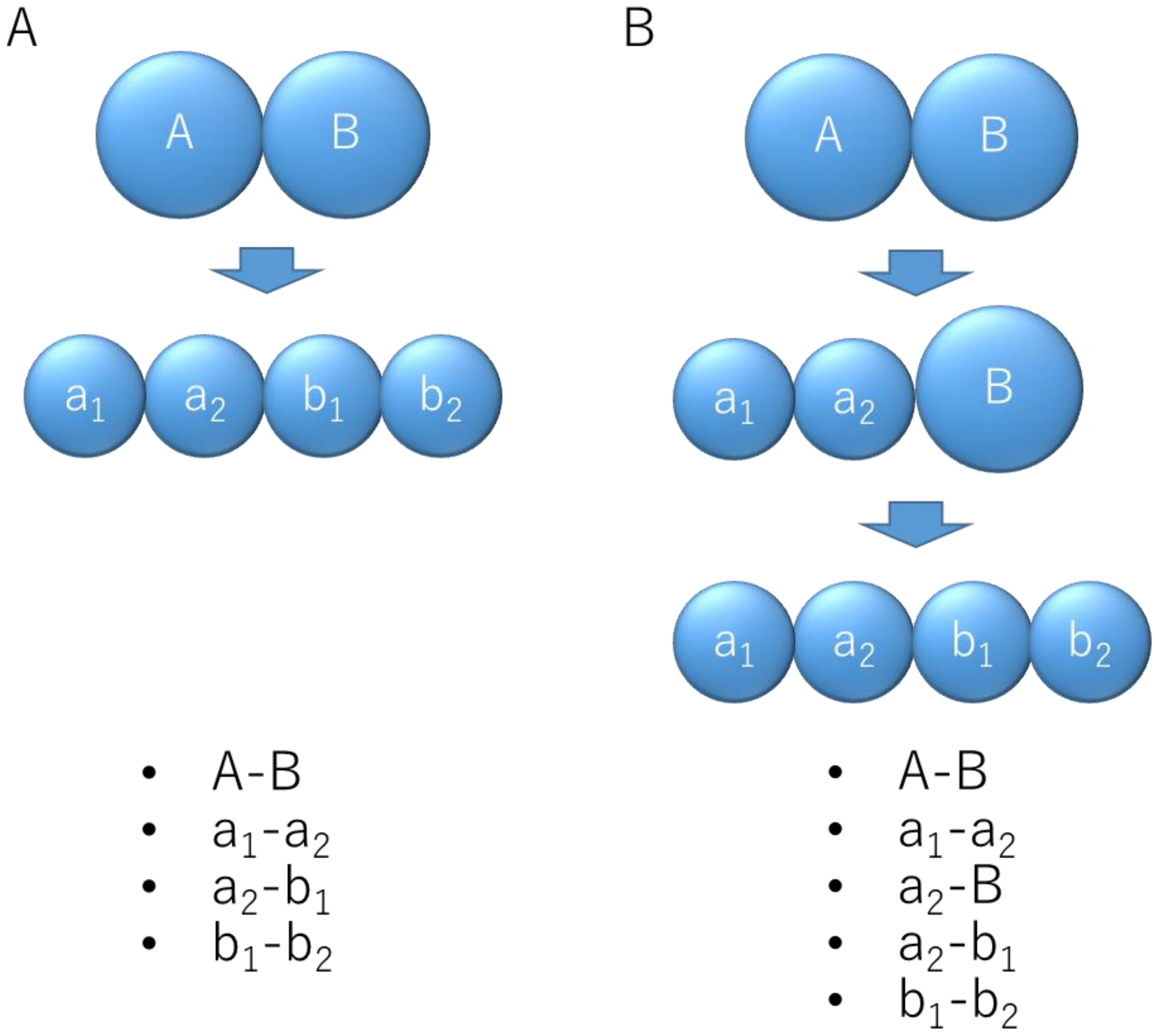


**Supplementary Figure 6.** Schematic explanation of variability of contacts caused by variability in division timings.

Schematics showing that differences in detected contacts are caused by variability in division timings. A and B are mother cells. a1 and a2 are daughter cells of A. b1 and b2 are daughter cells of B. (A) A and B divide concomitantly. (B) A divided one-timepoint faster than B. a2-B is additionally detected.

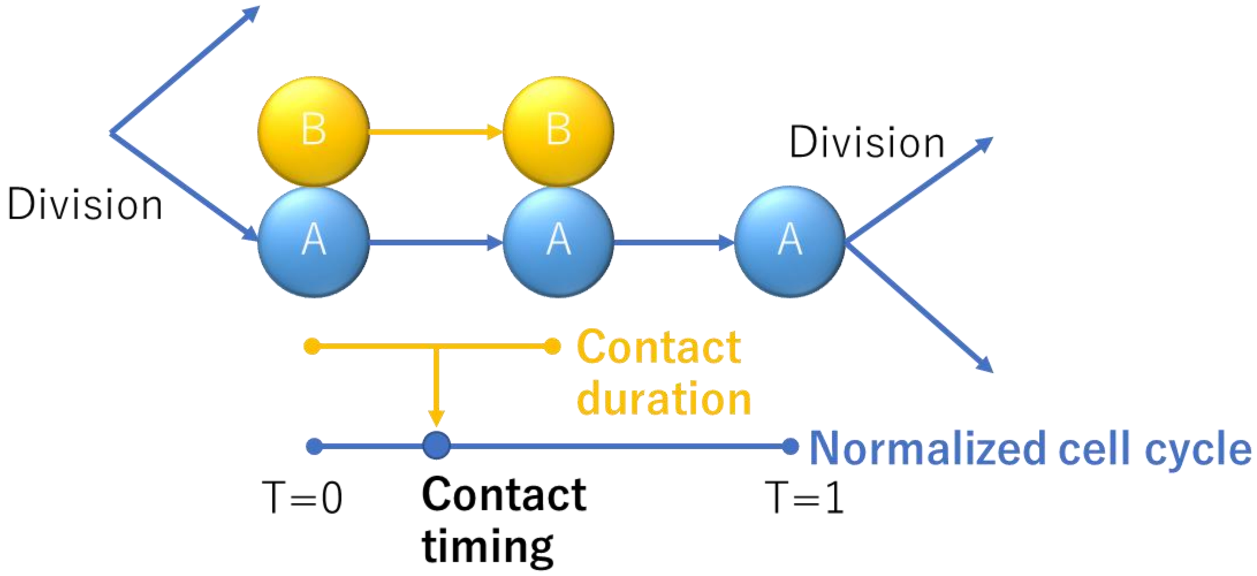


**Supplementary Figure 7.** Schematic explanation of the contact timing.

Schematics showing how to calculate the contact timing of A and B cells. It is the middle of contact duration within the normalized cell cycle period.

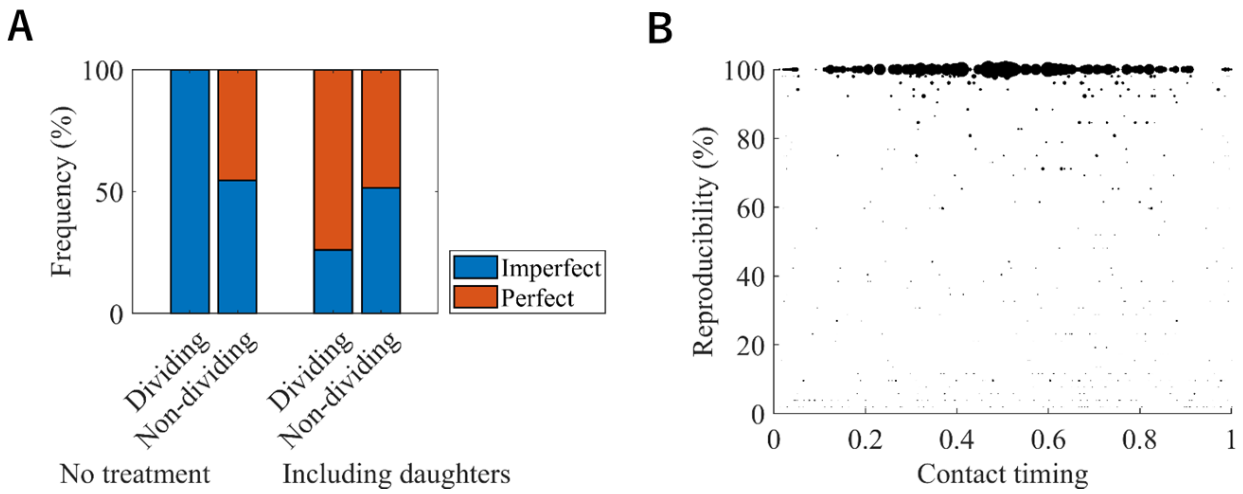


**Supplementary Figure 8.** Decrease of imperfectly reproducible contacts by removing variability in division timings.

(A) The proportion of perfectly and imperfectly reproducible contacts in dividing and non-dividing cells. The proportion of perfectly reproducible contacts increased by regarding contacts by their daughters as their contacts. (B) The reproducibility of the contact over the contact timing after removing variability caused by variability in division timings. Area indicates the integral area averaged for the cell and its daughters. The contacts with lower reproducibility on both sides observed in Figure 4B disappear. Instead, perfectly reproducible contacts are observed.

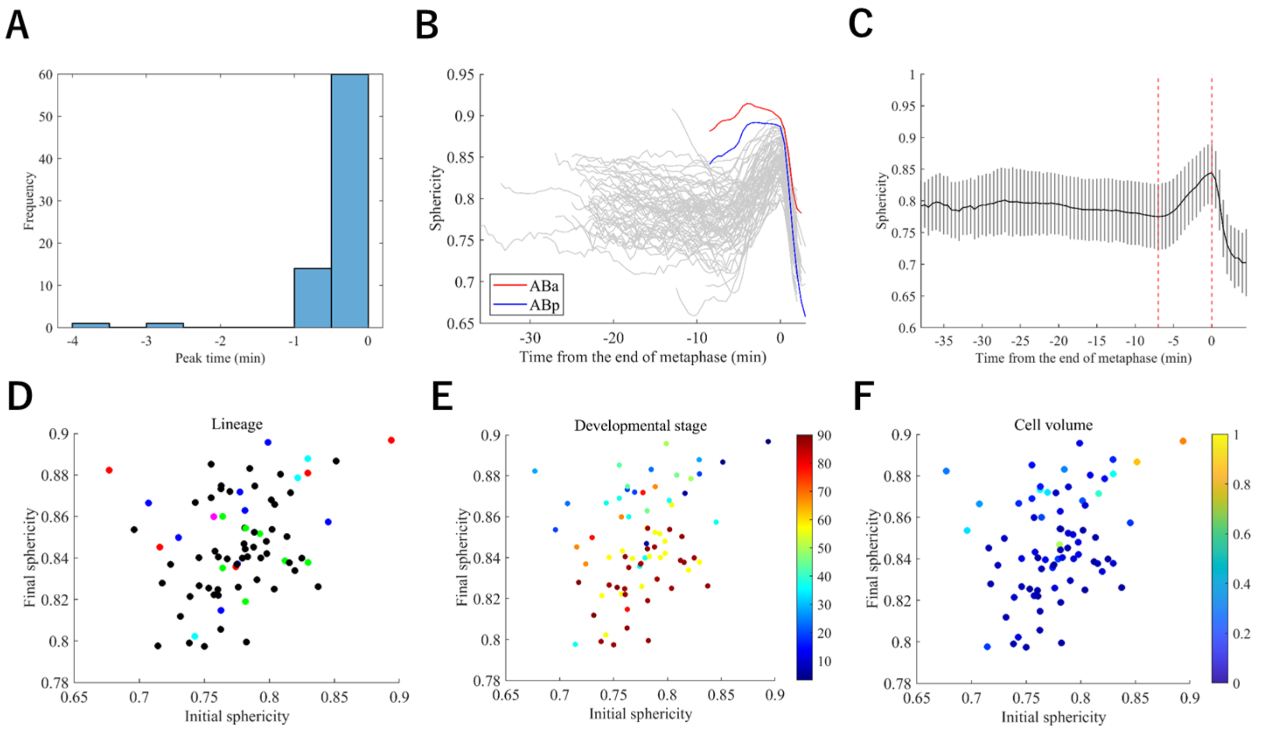


**Supplementary Figure 9.** Figures related to the mitotic rounding analysis.

(A) The distribution of sphericity peak times. (B) The sphericity dynamics of ABa and ABp. (C) The sphericity dynamics averaged over all cell types. Red vertical dotted lines indicate the initial and end of mitotic rounding. (D-F) Correlation between the initial and the final time points of the mitotic rounding colored by cell lineage (C), developmental stage (birth time) (E), and normalized cell volume (F).

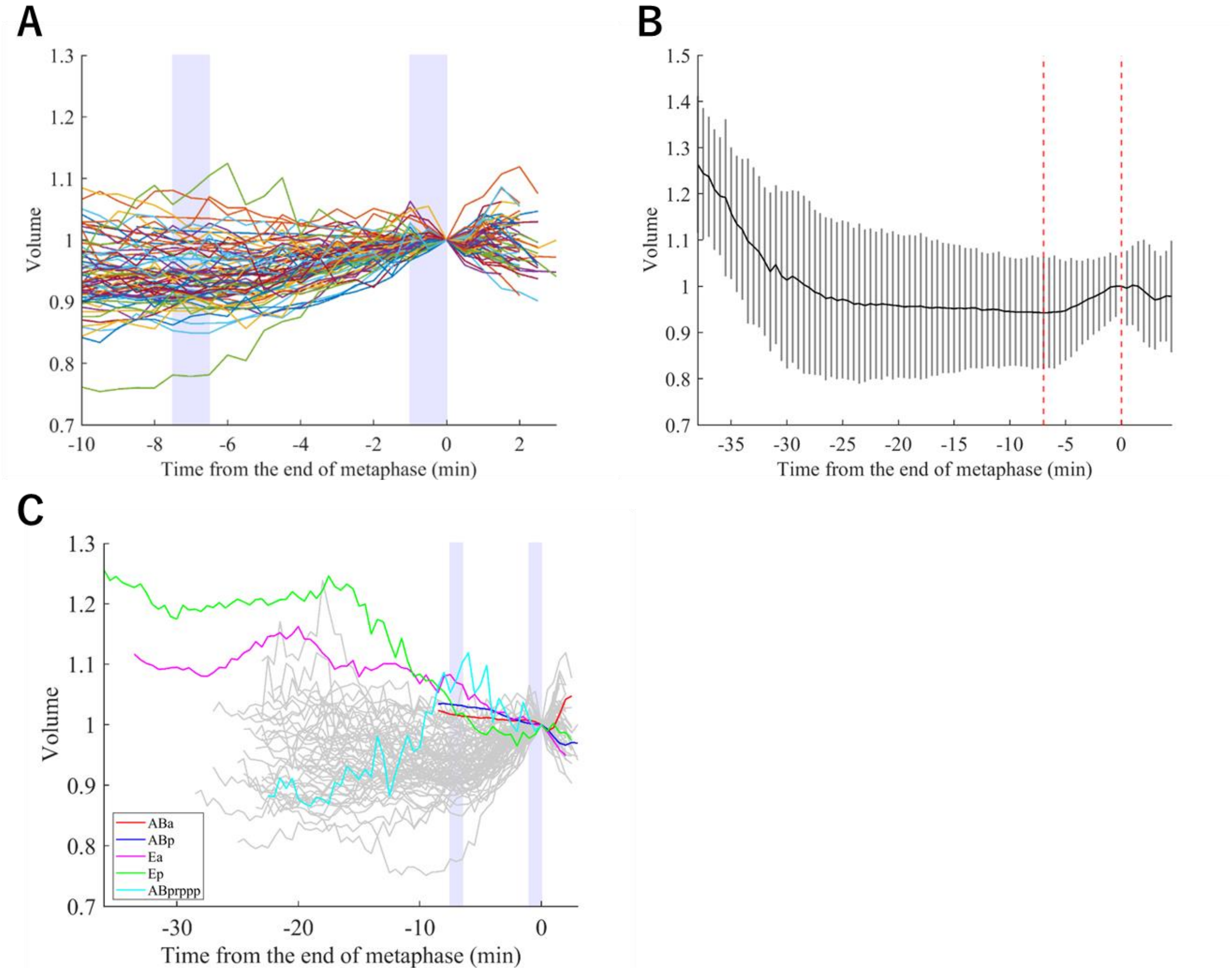


**Supplementary Figure 10.** Figures related to the mitotic swelling analysis.

(A) The cell volume dynamics around the period of the mitotic rounding. (B) The cell volume dynamics averaged over all cell types. Red vertical dotted lines indicate the initial and end of mitotic rounding. (C) The volume dynamics of cells that did not show the mitotic swelling.

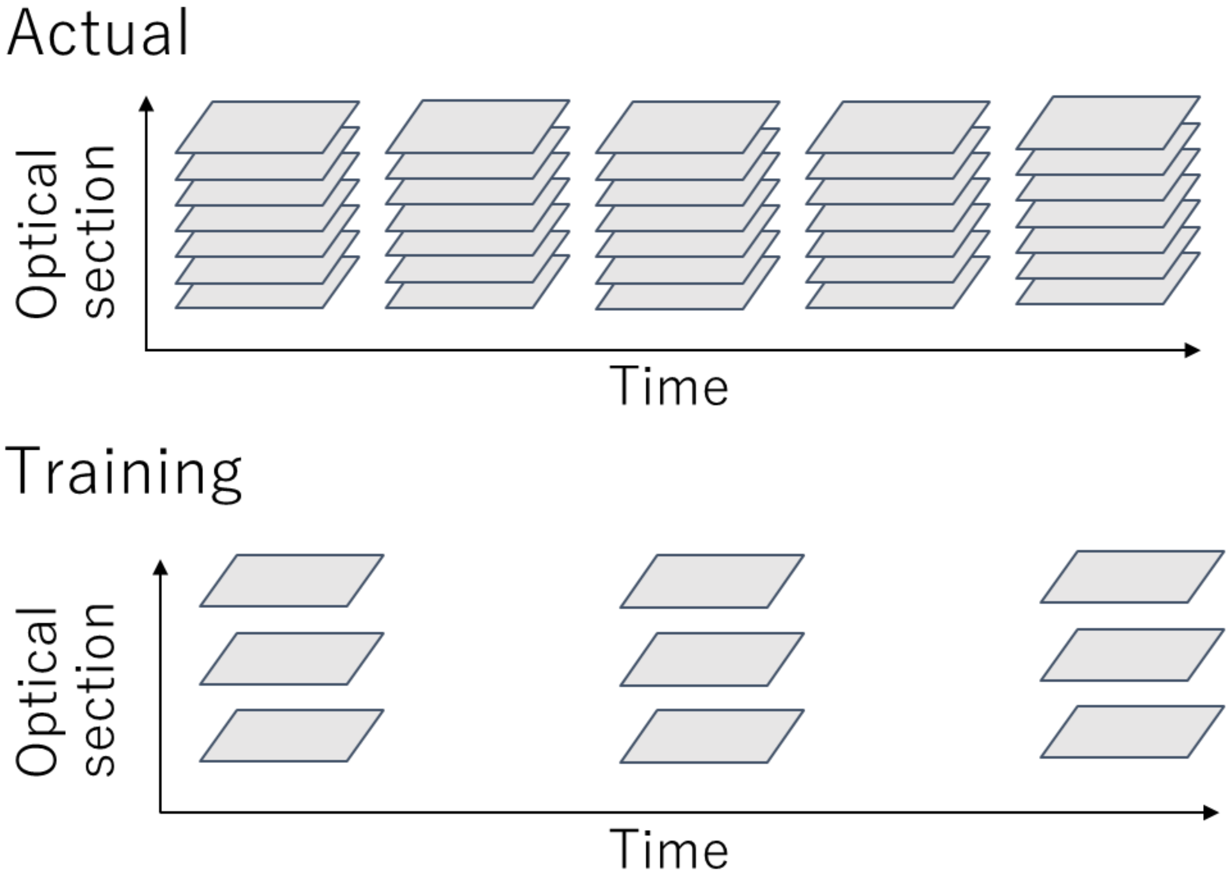


**Supplementary Figure 11.** Data sampling in the image restoration.

### 1.2 Supplementary Tables

**Supplementary Table 1.** Classification of the features.

| Category | Feature name | Details |
|---|---|---|
| Morphology | Volume | The number of voxels multiplied by the resolution |
| | Surface area | The number of voxels on the surface multiplied by the resolution |
| | SA/V | The surface area divided by the volume |
| | Solidity | The volume of the region divided by the volume of the smallest convex polygon that can contain the region |
| | Extent | The volume of the region divided by the volume of the smallest cuboid containing the region |
| | Sphericity | The extent to which the shape of the region resembles the perfect sphere |
| | Long axis | Length of the first PCA axis of the ellipsoid that has the same normalized second central moments as the region |
| | Middle axis | Length of the second PCA axis of the ellipsoid that has the same normalized second central moments as the region |
| | Short axis | Length of the third PCA axis of the ellipsoid that has the same normalized second central moments as the region |
| | Middle/Long | Middle axis divided by Long axis |
| | Short/Long | Short axis divided by Long axis |
| Cell-cell contact | Contact area | The number of voxels on the contact between the two cells |
| | Contact duration | The number of time points during which the contact is formed |

**Supplementary Table 2.** Cell-cell interactions used in this study and analysis results.

| Cell-cell interaction | Cell1 | Cell2 | Reproducibility | Integral area ($\mu m^2$) | Duration | Mean area ($\mu m^2$) | #Embryos | Reference |
|---|---|---|---|---|---|---|---|---|
| 1st Notch | ABp | P2 | 100 | 8842 | 23 | 385 | 52 | (Priess, 2005) |
| 2nd Notch | MS | ABalp | 100 | 3086 | 12 | 263 | 52 | (Priess, 2005) |
| 2nd Notch | MS | ABara | 100 | 1338 | 11 | 120 | 52 | (Priess, 2005) |
| 3rd Notch | ABplaaa | ABalapa | 100 | 4198 | 54 | 78 | 52 | (Priess, 2005; Chen et al., 2018) |
| 3rd Notch | ABplaaa | ABalapp | 100 | 3038 | 54 | 57 | 52 | (Priess, 2005; Chen et al., 2018) |
| 4th Notch | MSapa | ABplpapp | 100 | 782 | 17 | 44 | 32 | (Priess, 2005; Chen et al., 2018) |
| 4th Notch | MSapp | ABplpapp | 100 | 1073 | 18 | 63 | 32 | (Priess, 2005; Chen et al., 2018) |
| 5th Notch* | ABplpppp | MSappp | 100 | 91 | 2 | 46 | 2 | (Priess, 2005; Chen et al., 2018) |
| Wnt | EMS | P2 | 100 | 8077 | 29 | 276 | 52 | (Eisenmann, 2005) |
| Wnt | ABar | C | 100 | 1272 | 14 | 94 | 52 | (Walston et al., 2004) |

* The cell cycle does not complete

**Supplementary Table 3.** Manual annotation results of variable contacts.

| Cell1 | Cell2 | Integral area (μm$^2$) | Duration (time point) | Area(μm$^2$) | Annotation* |
|---|---|---|---|---|---|
| ABa | P2 | 12.8 | 1.3 | 11.2 | FP |
| ABar | ABplp | 32.5 | 1.2 | 26.1 | VCA |
| P3 | ABarp | 73.3 | 1.0 | 73.3 | FP |
| ABala | ABplap | 25.6 | 2.2 | 10.8 | VCA |
| ABalp | Ep | 40.3 | 1.0 | 40.3 | FP |
| ABara | ABalpp | 0.8 | 1.0 | 0.8 | VCA |
| ABarp | MSa | 333.9 | 16.0 | 17.7 | VCA |
| ABpra | ABplap | 4.8 | 2.0 | 2.4 | VCA |
| ABprp | ABarpp | 44.1 | 2.0 | 17.2 | VCA |
| MSa | ABprpa | 39.4 | 1.5 | 30.1 | FP |
| MSp | D | 20.8 | 1.0 | 20.8 | FP |
| Ea | Cp | 482.4 | 29.7 | 12.1 | VCA |
| Ea | ABaraa | 28.0 | 1.5 | 17.2 | FP |
| Ea | ABarpp | 514.1 | 16.7 | 24.6 | VCA |
| Ca | MSap | 127.7 | 9.8 | 11.4 | VCA |
| ABplap | ABprpa | 10.8 | 1.0 | 10.8 | FP |
| ABalaa | ABarap | 49.9 | 1.8 | 23.0 | FP |
| ABalpa | MSap | 354.3 | 10.8 | 38.8 | VCA |
| ABalpp | ABarpa | 68.3 | 5.6 | 10.0 | VCA |
| ABarpa | MSaa | 258.3 | 12.3 | 15.8 | VCA |
| ABarpp | MSpp | 600.5 | 24.5 | 19.8 | VCA |
| ABprpa | MSap | 53.6 | 2.6 | 14.9 | FP |
| ABprpp | MSpa | 27.8 | 2.0 | 14.8 | FP |

*VCA: variability in cell arrangement, FP: false positive

**Supplementary Table 4.** Features used for collection of the time lag.

| Feature name | Definition |
|---|---|
| Time | Time after nuclear division |
| Membrane intensity | Maximum intensity on the line connecting centers of the nuclei in membrane image |
| Membrane intensity ratio | The ratio of Membrane intensity to the previous time point |
| Membrane intensity thus far | Maximum Membrane intensity until this time point |
| Position of max intensity | The position of Max membrane intensity on the line connecting centers of the nuclei |
| Nuclear intensity | Mean intensity of the nuclei |
| Nuclear intensity ratio | The ratio of Nuclear intensity to the previous time point |
| Nuclear size | Total size (number of pixels) of the nuclei |
| Nuclear size ratio | The ratio of Nuclear size to the previous time point |
| Nuclear distance | Distance between the centers of nuclei |
| Nuclear distance ratio | The ratio of Nuclear distance to the previous time point |

### 2 Supplementary videos

**Supplementary Video 1.** Segmentation results of cells that completed the cell cycle in an embryo.

**Supplementary Video 2.** The dynamics of ABplap and ABalpp.

The cellular regions of ABplap and ABalpp are colored in red. The video shows both cells do not contact.

**Supplementary Video 3.** The dynamics of ABplap and ABplpp.

The cellular regions of ABplap and ABplpp are colored in red. The video shows both cells do not contact.
